# Temporal synchrony between human odor rhythms and mosquito olfactory preference shapes host attraction

**DOI:** 10.64898/2026.01.18.700155

**Authors:** Lan Lou, Karthikeyan Chandrasegaran, Julien Devilliers, Austin Compton, Abdulhadi Kobiowu, Emilie Applebach, Adaline Bisese, Richard Rust, Shajaesza Diggs, Sydney Luff, Sneha Sapkota, Nicole E. Wynne, Diane F. Eilerts, Olivia Evans, Joshua B. Benoit, Zhijian Jake Tu, Chloé Lahondère, Clément Vinauger

**Affiliations:** Department of Biochemistry, Virginia Polytechnic Institute and State University, Blacksburg, VA 24061, USA; Department of Entomology, University of California Riverside, Riverside, CA 92521, USA; Genetics, Bioinformatics and Computational Biology (GBCB), Virginia Polytechnic Institute and State University, Blacksburg, VA 24061, USA; Department of Biological Sciences, University of Cincinnati, Cincinnati, Ohio 45221, USA; Center for Emerging Zoonotic and Arthropod-borne Pathogens, Blacksburg, VA 24061, USA; Fralin Life Science Institute, Blacksburg, VA 24061, USA; The Global Change Center, Virginia Polytechnic Institute and State University, Blacksburg, VA, USA; Center for the Mathematics of Biosystems, Virginia Polytechnic Institute and State University, Blacksburg, VA 24061, USA

**Keywords:** Circadian rhythms, Olfactory sensitivity, Host-seeking behavior, Temporal synchronization, *Aedes aegypti*, Human body odor, Chronobiology, Vector-host interactions

## Abstract

For anthropophilic mosquitoes such as *Aedes aegypti*, aligning host-seeking with human availability enhances foraging efficiency and reproductive success. Although time of day modulates mosquito activity and olfactory sensitivity, it remains unknown whether human hosts display rhythmic changes in odor cues and whether mosquitoes adjust their sensory responses accordingly. Here, we combine chemical, behavioral, genetic, and transcriptomic approaches to reveal that both mosquitoes and their human hosts in this interaction are temporally synchronized. Gas chromatography–mass spectrometry showed systematic daily shifts in human body odor composition between morning and evening. Correspondingly, mosquitoes prefer host odors that match their own active phase—a time-specific preference abolished in *timeless* mutants and under constant darkness. Silencing the *timeless* gene further induced an aversion for the host scent under light-dark conditions. Transcriptomic analysis of mosquito heads and antennae uncovered rhythmic expression of sensory and neuromodulatory genes, driven by both circadian and light–dark cycles and which peaks during mosquitoes’ active periods, with rhythmic co-expression networks collapsing in timeless knockouts. Together, these results show that mosquito attraction to humans is temporally tuned by the interplay of host odor rhythms and mosquito sensory rhythms, revealing a previously unrecognized form of interspecific temporal synchronization in vector-host interactions.

## Introduction

Daily rhythms are ubiquitous, allowing organisms to anticipate and adapt to predictable environmental changes across the day-night cycle (1, 2). By synchronizing physiology and behavior with these temporal variations, plants and animals optimize resource use and survival (2, 3). In addition, interactions between organisms are often rhythmic as well: prey avoid predators, pollinators visit flowers, and mates locate each other at specific times of day (4–6). Such temporal coordination can arise through environmentally driven diel cycles and the action of endogenous circadian clocks, aligning behavioral and physiological processes of interacting species to maximize mutual benefit or, in antagonistic contexts, to confer competitive advantage.

For disease-vector arthropods, aligning host-seeking activity with periods of maximal host availability provides a clear fitness benefit (4, 7, 8). Nocturnal disease vectors, such as Anopheline mosquitoes and triatomine bugs feed primarily when human hosts are asleep, immobile, and consequently, less defensive (9). These species can even shift their activity rhythms in response to changes in host behavior or accessibility, demonstrating a high degree of temporal plasticity (10–12). In contrast, diurnal mosquitoes like the yellow fever mosquito *Aedes aegypti* interact with active, mobile hosts (13), which imposes distinct selective pressures on host-seeking behavior, likely favoring temporal flexibility to optimize chances of getting a blood meal and surviving host interactions. Their host-seeking behavior typically peaks during crepuscular periods, early morning and late afternoon, when humans are most available but less defensive (14–17). This timing reflects an adaptive balance between physiological readiness for a potential blood meal and reduced predation risk (4, 18–20).

Temporal organization in mosquitoes is underpinned by rhythmic gene expression across multiple timescales, which coordinate development and daily behaviors. Across a single day, diel rhythms in locomotor activity, host-seeking, and olfactory sensitivity (14, 21) correspond to rhythmic expression of sensory genes (21–23). Microarray studies indicate that 6-20 % of *Ae. aegypti* head transcripts and roughly 15 % in *Anopheles gambiae*, exhibit diel or circadian regulation (21, 22, 24). Interestingly, in both species, transcriptional activity peaks prior to respective activity times, just before behavioral activity onset, suggesting anticipatory regulation that primes mosquitoes for host-seeking (21).

Human body odor, a complex blend of over 400 volatile compounds (25, 26), is a key cue used by *Ae. aegypti* to locate hosts (27, 28). This species detects and discriminates specific components such as sulcatone, lactic acid, and carboxylic acids to distinguish humans from other animals and to select preferred individual hosts (28–32). While the rhythmicity of mosquito host-seeking behavior is well characterized, the daily changes in host odor composition and whether mosquito sensory systems are adaptively tuned to daily temporal variations remain unknown.

Here, we demonstrate that human body odor fluctuates across the day, and that *Ae. aegypti* females modulate their olfactory preferences in a time-dependent manner controlled by environmental cues and maintained by the circadian clock. Combining behavioral assays using time-specific synthetic odor blends, transcriptomic profiling of both the head and antennae, and a newly generated *timeless* knockout line, we reveal that clock-independent modulation of olfactory sensitivity synchronizes mosquito host-seeking preferences with the temporal dynamics of human odor production. These findings uncover a previously unrecognized dimension of mosquito–host interaction, in which rhythms in sensory processes align vector attraction with rhythmic host cues, enhancing temporal precision in host detection. Because host-seeking underlies biting and pathogen acquisition, this temporal tuning may also influence when transmission risk is higher and interventions are more effective.

## Results

### Human body odor varies throughout the day

*Aedes aegypti* females exhibit two daily peaks in spontaneous locomotor activity—a minor one at dawn and a more pronounced one at dusk (Fig. 1A). This bimodal pattern has been consistently documented across laboratory and field studies (14, 15, 33). Previous work has examined the specific behaviors performed during these active phases. While sugar feeding can occur throughout the day, host-seeking and blood-feeding are typically concentrated around the crepuscular periods, with stronger responses to host cues at dusk (34) and occasional increases in morning biting during the dry season (17). Overall, *Ae. aegypti* tends to bite hosts preferentially during its evening activity peak.

**Fig. 1.**
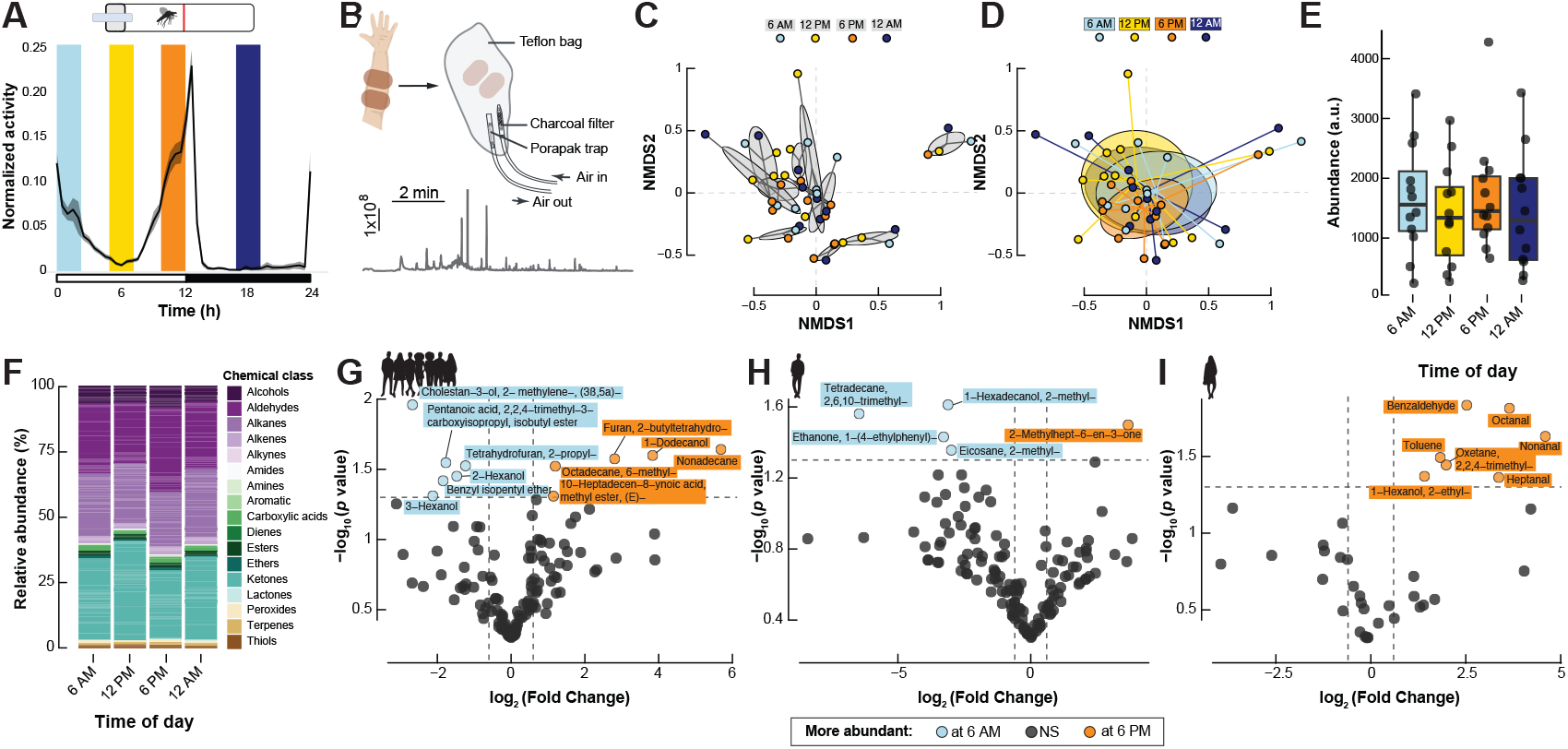
Human body odor profiles exhibit subtle yet consistent chemical shifts which align with mosquito host-seeking daily rhythms. **(A)** Schematic diagram of the mosquito locomotor activity monitoring system (upper) and activity pattern for *Aedes aegypti* females in a 12:12 hr light-dark (LD) cycle (lower). The x-axis indicates *Zeitgeber* time (ZT), with white and black bars denoting light and dark phases, respectively. The shaded area around the trace represents the standard error of the mean (n = 100). The colored vertical bars indicate the four ZT time points selected for subsequent analysis: two corresponding to peak activity (dawn and dusk) and two representing trough periods. **(B)** Schematic of the host odor collection setup, in which human-worn nylon sleeves were enclosed in a Teflon bag with a charcoal-filtered air stream. Volatiles were trapped on a Porapak filter for subsequent analysis. A representative GC-MS chromatogram of human odor samples is shown below. **(C)** Non-metric multidimensional scaling (NMDS) of human odor samples grouped by host identity. Odor profiles differed significantly among individuals (PERMANOVA, *p* = 0.001). **(D)** NMDS of human odor samples grouped by sampling time (6 AM, 12 PM, 6 PM, 12 AM). No significant clustering was observed across time points (PERMANOVA, *p* = 0.998; n = 44). **(E)** Total abundance of human body odor volatiles throughout the day. **(F)** Relative abundance of chemical classes in human body odor profiles across time of day. Odor samples were pooled by time points before analysis. **(G-I)** Volcano plots showing differentially enriched compounds at 6 AM *vs*. 6 PM in **(G)** a combined multi-host dataset, **(H)** a representative male host and **(I)** a representative female host. Compounds significantly more abundant at 6 AM are shown in blue, and those enriched at 6 PM in orange (log_2_ (fold-change) > 0.6, adjusted *p* < 0.05).

To test whether this crepuscular biting pattern is correlated with daily changes in the composition or abundance of human odor cues, we collected body odor samples from eleven volunteers at four time points corresponding to the mosquitoes’ activity peaks (6 AM and 6 PM) and troughs (12 PM and 12 AM) (Fig. 1A). Volatiles emitted from nylon sleeves worn on the forearms were analyzed by gas chromatography–mass spectrometry (GC-MS) (Fig. 1B). In total, 316 odor samples were collected, including clean-sleeve controls, with at least four independent replicates per volunteer-time combination, each collected on a separate day to control for intra-individual variation.

A total of 1239 distinct volatile compounds across all samples were identified by GC-MS. To reduce noise from trace compounds that were both low in abundance and inconsistently detected (31), we removed chemicals representing <1 % of the total abundance in each sample, retaining 590 compounds for further analysis. To characterize odor variation among multiple hosts, we focused on compounds detected in at least 7 of the 11 individuals and/or previously reported as key markers of human body odor, yielding a final set of 362 volatiles (see full dataset for details).

As expected, human odor profiles varied substantially among individuals (PERMANOVA, *p* = 0.001; Fig. 1C). Although time of day did not significantly influence the overall chemical composition (PERMANOVA, *p* = 0.998 at the group level; *p* > 0.332 at the individual levels; Fig. 1D), individual odor signatures showed measurable within-day variation (0.04 < s.d. < 0.27; Fig. 1C). Notably, samples collected at 6 PM showed a subtle convergence toward a more similar chemical profile across participants (mean dispersion 0.25 s.d. *vs*. 0.29–0.37 at other time points; ANOVA, *p* = 0.56; Fig. 1D). This suggests that human odor composition may be most stereotyped in the early evening, coinciding with the primary host-seeking period of *Ae. aegypti*.

The subtle daily variations in human odor profiles could reflect either quantitative or qualitative changes in scent composition. Total volatile abundance varied significantly among individuals (Generalized Linear Model (GLM), *p* < 0.001; Fig. 1E) but not across sampling times (GLM, *p* = 0.83; Fig. 1E). Thus, overall scent intensity remained stable throughout the day, suggesting that enhanced mosquito attraction in the evening is not driven by changes in odor quantity.

We next examined qualitative variation in odor composition by grouping volatiles into functional classes (Fig. 1F, Fig. S1). Human scent was dominated by ketones (daily average: 28.2 ± 2.16 %), aldehydes (27.55 ± 2.18 %), alkanes (19.86 ± 0.29 %), and alcohols (9.69 ± 0.21 %), consistent with previous reports (20, 25, 31, 35). Carboxylic acids, known contributors to individual host attractiveness (32), represented only 2.09 ± 0.13 % of total volatiles, likely reflecting the lower recovery of this chemical class under our sampling conditions (36). The relative proportions of chemical classes did not significantly vary over time (GLMM, *p* = 0.615), indicating that the overall functional composition of human odor remained stable throughout the day.

Despite this apparent stability, mosquito olfaction is highly sensitive to subtle shifts in volatile ratios. Even minor changes in compounds such as sulcatone, aldehydes, or carboxylic acids can strongly influence attraction (29, 32, 37). To identify such fine-scale time-dependent variation, we compared compound abundances between pairs of sampling time points (Fig. S1). At the group level, eleven volatiles were significantly differentially enriched between odors collected at 06:00 and 18:00 (Fig. 1G). These compounds emerged as consistent signals despite considerable interindividual and day-to-day variability, suggesting systematic daily changes in human scent composition.

At the individual level, volunteers each displayed compounds that differed significantly between 6 AM and 6 PM (see Fig. 1H,I for representative male and female volunteers). Compared to pooled analyses, individual odor profiles were more internally consistent (Fig. 1D), enabling clearer detection of temporally modulated volatiles that might otherwise be masked by inter-individual variation. Additional pairwise comparisons among other time points (6 PM *vs*. 12 PM, 6 PM *vs*. 12 AM, and 12 PM *vs*. 12 AM) revealed further time-specific differences in compound abundance (Fig. S1C-E). Collectively, these results indicate that the human odor composition undergoes subtle yet reproducible diel fluctuations, particularly between morning and evening, which may provide mosquitoes with reliable temporal cues for host seeking.

### Mosquito preferences and host scent rhythms are phase-aligned

Based on the differently enriched compound analysis (Fig. 1G-I, Fig. S2) and knowledge on chemicals with well-established roles in mosquito attraction from the literature (29, 38–40), we selected twelve key compounds (nonanal, octanal, 2-hexanone, heptanal, 2-hexanol, 1-hexadecanol, glutaric acid, nonanoic acid, hexanal, propenoic acid, pentanal and sulcatone) for the preparation of an artificial human odor blend. These compounds were chosen because they consistently varied across time points and, together, captured the core chemical features of human scent profiles relevant to mosquito attraction. Using data from a representative male volunteer, for whom additional replicates (n = 74) were conducted to ensure robustness and account for day-to-day variation, we prepared time-specific artificial blends recapitulating the relative abundances of the selected compounds at either 6 AM or 6 PM (referred to as the “AM blend” and the “PM blend”, respectively). To optimize the artificial blends for behavioral testing, we adjusted concentrations according to each compound’s vapor pressure, enabling accurate replication of natural chemical release dynamics under experimental conditions (Fig. 2A,B).

**Fig. 2.**
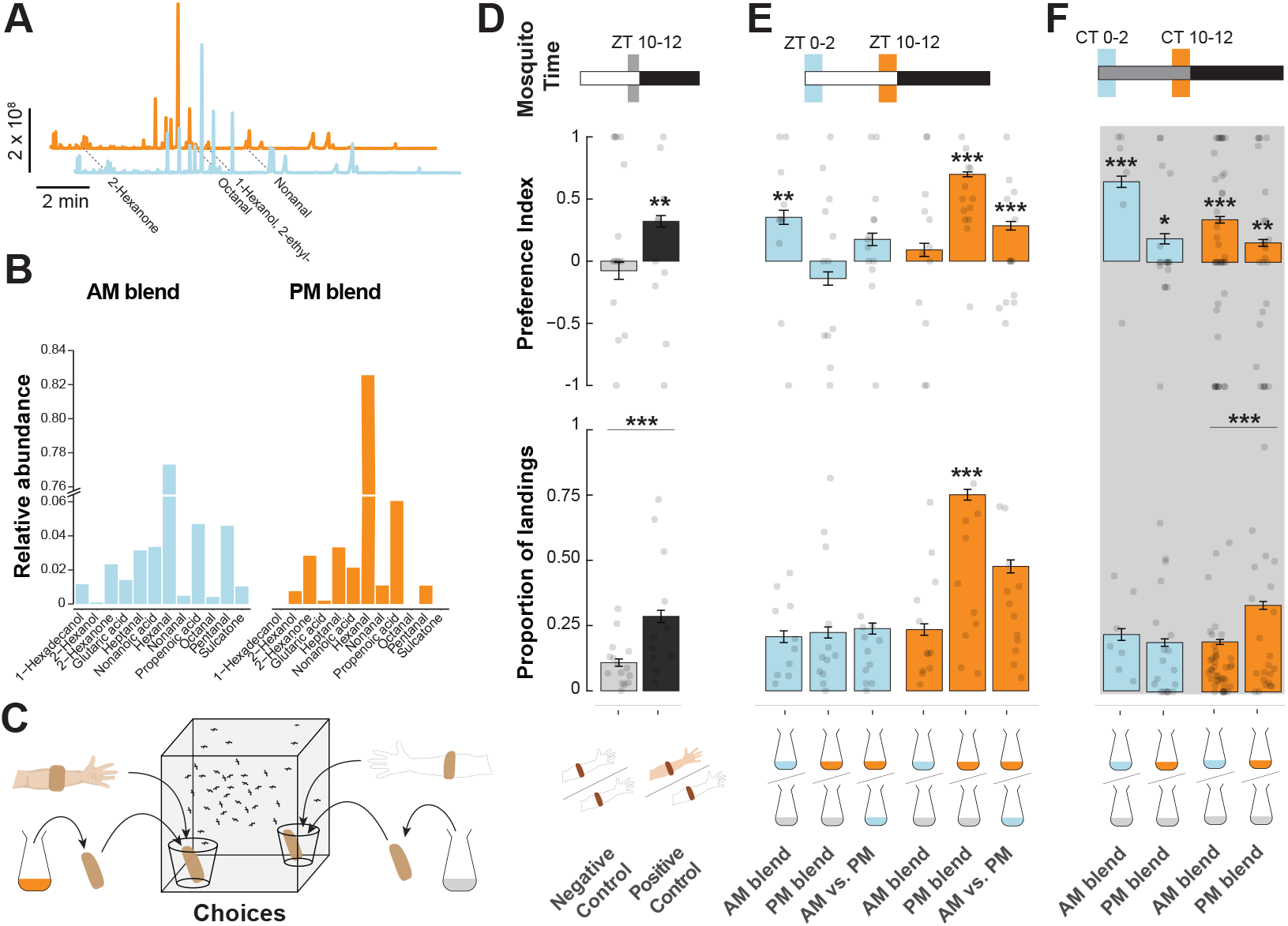
Light cues drive the temporal alignment between host odor rhythms and mosquito olfactory preference. **(A)** Representative chromatograms highlighting a selection of the key chemical compounds of the AM (light blue) and PM (orange) blends. **(B)** Relative composition of artificial human odor blends mimicking scents collected at 6 AM (left, blue) and 6 PM (right, orange). **(C)** Schematic of the dual-choice landing assay used to quantify mosquito attraction. **(D-F)** Adult female *Aedes aegypti* were tested in a dual-choice landing assay, where they could select between either two sleeves contained in meshed plastic cups. Mosquitoes were released into a cage and allowed to land on either option (see lower left for setup). The top panel shows the preference index based on landing counts. The bottom panel visualizes the proportion of mosquitoes that landed on either sleeve. **(D)** Control mosquitoes given the choice between two clean sleeves (light gray, neutral control) or a human-worn sleeve versus a clean one (positive controls). **(E)** Behavioral preference and landing proportion under light–dark conditions, at ZT 0-2 (light blue) or ZT 10-12 (orange) when mosquitoes were offered odor blends mimicking 6 AM (light blue), 6 PM (orange), or control (gray) odors. **(F)** Behavioral preference and landing proportion under constant darkness at CT 0-2 (light blue) or CT 10-12 (orange) for the same odor blends as in (E). Asterisks in top panels D, E and F denote statistically significant preference relative to random choice (Binomial Exact Test, *p* < 0.05). Asterisks in the bottom panel D indicate significant difference between the positive and neutral controls. Asterisks in the bottom panels E and F indicate a significant difference with the positive control (Tukey post hoc test, * *p* < 0.05; ** *p* < 0.01; *** *p* < 0.001).

To test whether these temporal variations in human odor influence mosquito attraction, we used a dual-choice landing assay (Fig. 2C). A total of 11,323 mosquitoes were tested, distributed in 295 groups of 13 < N < 73 individuals (38 ± 10 individuals per group), across 20 experiments (see raw data for details). Among them, 2,383 mosquitoes, or 21 % of the released individuals, chose either of two nylon sleeves enclosed in mesh-covered plastic cups.

When quantifying the preference between two clean sleeves (*i*.*e*., odorless), mosquito landing activity was low (10.8 ± 1.4 %; n = 15, N = 482). The corresponding preference index (PI = -0.08 ± 0.06) was not different from zero, *i*.*e*., an equal distribution of landings between the two sleeves (Binomial Exact test: *p* = 0.67), indicating no directional bias. In contrast, when one sleeve had been worn by a human volunteer (n = 11, N = 372), mosquitoes showed a significant preference for the odor-treated sleeve (PI = 0.32 ± 0.05, *p* = 0.0012) and the overall responsiveness significantly increased (landing proportion = 28.5 ± 2.3 %, *p* < 0.001; Fig. 2D).

After verifying that human-worn sleeves showed distinct increases in mosquito landing, we shifted our focus to exam-ining how the AM and PM blends impact landing. Overall, the type of artificial blend presented, the time of the day, and the light regime (*i*.*e*., light-dark or constant darkness) were significant predictors of the mosquito preferences (GLM, *p* < 0.031). Importantly, the use of the PM blend under light-dark conditions was a strong predictor of mosquito preference (GLM, *p* < 0.001) and the interaction between the PM blend, the morning time, and a light-dark regime was also significant (*p* = 0.002). Similarly, the use of the PM blend was a significant predictor of the proportion of mosquitoes landing on either cup (GLM, *p* < 0.001), with the interaction between the use of the PM blend, light-dark conditions, and time of day being significant (*p* = 0.002).

Under light-dark (LD) conditions (Fig. 2E), mosquitoes tested at *Zeitgeber* Time (ZT) 0-2 showed a significant preference for the AM blend over an odorless control (PI = 0.35 ± 0.05, Binomial Exact test: *p* = 0.005, n = 10), and the over-all landing activity was comparable to the positive control in which mosquitoes were given the choice between a clean sleeve and a human host-worn sleeve (20.7 ± 2.24 %, Tukey-adjusted *p* = 0.796). No significant preference was observed for the PM blend versus control for ZT 0-2 mosquitoes (PI =-0.14 ± 0.05, Binomial Exact test: *p* = 0.235, n = 12), and landing activity was not different from the control (22.3 ± 2.12 %, Tukey-adjusted *p* = 0.941). When ZT 0-2 mosquitoes were given a choice between the AM and PM blends, no significant preference was detected (PI = 0.18 ± 0.04, Binomial Exact test: *p* = 0.104, n = 13; landing proportion = 23.8 ± 2.1 %, Tukey-adjusted *p* > 0.99).

At ZT 10-12 (Fig.2E), no significant preference was detected when mosquitoes were tested with the AM blend (PI = 0.09 ± 0.05, Binomial Exact test: *p* = 0.45, n = 12; landing proportion = 23.5 ± 2.2 %, *p* = 0.98). In contrast, mosquitoes showed a strong preference for the PM blend (PI = 0.70, Binomial Exact test: *p* < 0.001, n = 12), associated with a marked increase in landing activity (75.1 ± 2.1 %, Tukey-adjusted *p* < 0.001). When offered a choice between the AM and PM blends, ZT 10-12 mosquitoes preferred the PM blend (PI = 0.29 ± 0.03, Binomial Exact test: *p* < 0.001, n = 13), but their preference for the PM blend was lower than when the PM blend is presented alone (Tukey-adjusted *p* < 0.001). Although the landing proportion (47.7 ± 2.5 %) was lower than when the PM blend was presented alone, the difference was not significant and not different from the positive control (Tukey-adjusted *p* > 0.9 in both cases). Together, these results demonstrate that under LD conditions, mosquitoes preferentially respond to host odors that are temporally aligned with their own active phase, with a significant effect of the PM blend in the evening time on the number of mosquitoes making a choice.

Under constant darkness (DD) conditions (Fig. 2F), the temporal pattern of behavioral preferences was distinctively different from under LD (GLM, *p* = 0.031). Mosquitoes were significantly attracted by either blend at either time of day (Morning time [CT 0-2]: PI = 0.64 ± 0.04, Binomial Exact test: *p* = 0.004, n = 7, and PI = 0.18 ± 0.04, *p* = 0.029, n = 19for the AM and PM blends, respectively; Evening time [CT 10-12]: PI = 0.34 ± 0.02, *p* = 0.001, n = 42 and PI = 0.15 ± 0.03, *p* = 0.005, n = 24, for the AM and PM blends, respectively). At both time points, the preference for the AM blend tended to be more pronounced than for the PM blend but the difference was not statistically significant (Tukey-adjusted *p* = 0.057 in the morning and *p* = 0.38 in the evening time). At CT 0-2, the proportion of mosquitoes making a choice was not significantly different from the positive control when the either blend was used (Tukey-adjusted *p* = 0.89 and *p* = 0.14, for the AM and PM blend, respectively). At CT 10-12, when compared to the positive control, slightly fewer mosquitoes made a choice in the presence of the AM blend (*p* < 0.069) while slightly more mosquitoes made a choice when the PM blend was used (*p* = 0.99). When compared to each other, the proportion of mosquitoes making a choice was not significantly different between the two blends in the morning (Tukey-adjusted *p* = 0.99) but was significantly larger when the PM blend was used in the evening than when the AM blend was used (Tukey-adjusted *p* < 0.001).

Together, these results indicate a phase-dependent attraction to human odor cues driven by environmental light cues. Indeed, under LD, peak responsiveness occurred when the odor blend matched the evening profile of the host during the evening of the mosquitoes. Under constant darkness, this phase alignment was suppressed, with a tendency for a stronger preference for the AM blend, suggesting a baseline olfactory sensitivity tuned for the specific chemical ratios in this blend.

### Timeless disruption abolishes phase-aligned mosquito preferences

To investigate the role of the endogenous circadian clock in regulating the olfactory preference of mosquitoes, we generated a 91-bp deletion in the *timeless* gene using CRISPR/Cas9 (Fig. 3A, Fig. S3; see Methods for details). Knocking out *timeless* prevents the circadian molecular clock from entraining to external light cues by preventing the formation of PER:TIM heterodimers (proteins of *period* and *timeless* genes, respectively)(41– 43). The *timeless* mutants behaved similarly to wild-type individuals under light/dark conditions, although with a less defined morning and evening peaks and a slightly shallower trough in activity at midday, without obvious morphological or developmental differences compared to wild type mosquitoes. However, under constant darkness conditions, knocking out *timeless* resulted in an inability to maintain rhythmic locomotor activity patterns (Fig. 3B).

**Fig. 3.**
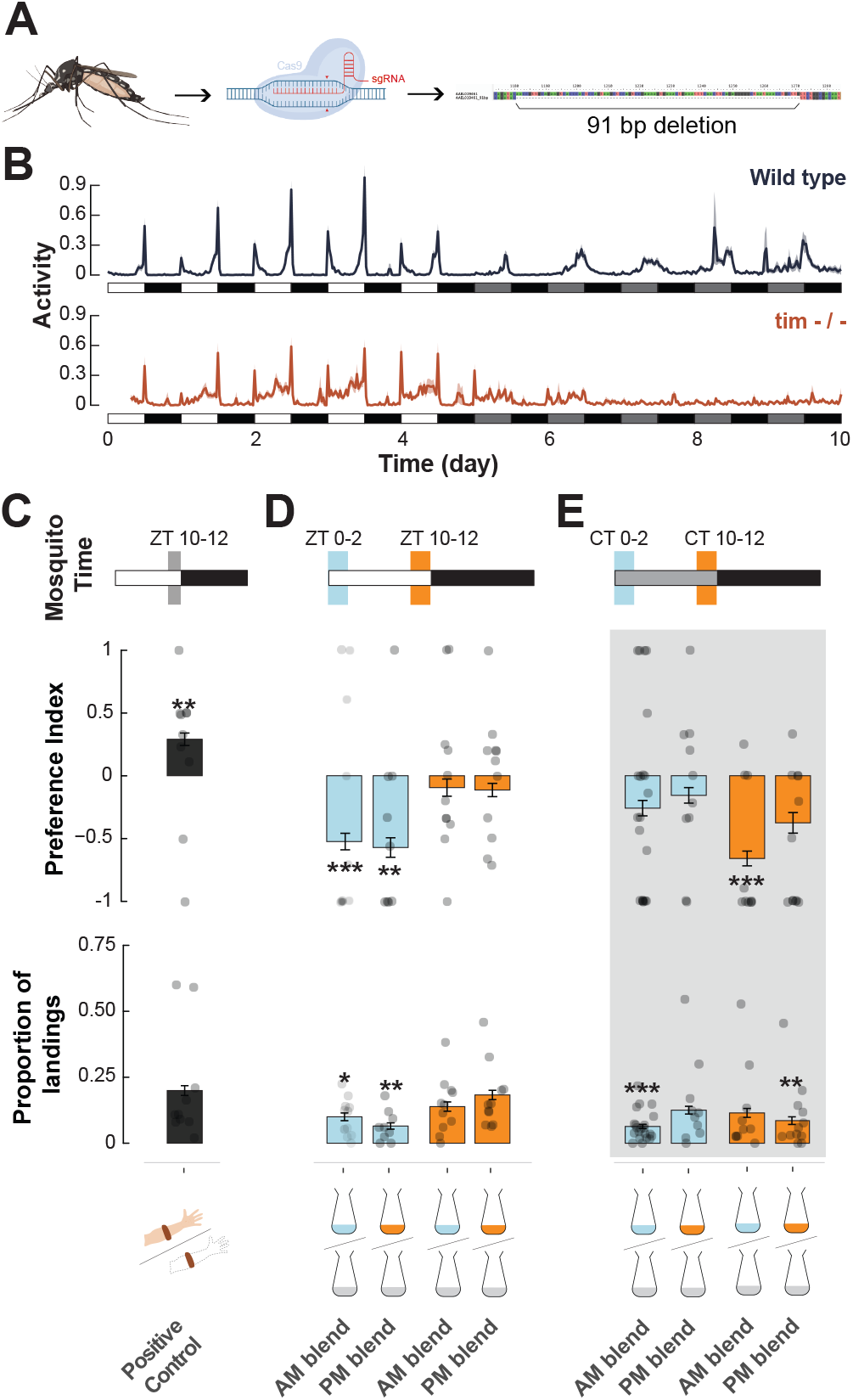
The gene *timeless* is required for the attraction to artificial blends, but responses to real hosts are *timeless*-independent. **(A)** Schematic representation of the generation of the *timeless* mutant line. A 91-bp deletion was introduced into the *timeless* coding region to disrupt gene function. **(B)** Locomotor activity pattern of the *timeless* mutant female *Aedes aegypti* under a 12:12 hr light-dark (LD) cycle and under a constant darkness (DD) condition (n = 55 for *timeless* and n = 88 for *wt*). The x-axis indicates *Zeitgeber* time (ZT) for LD, or circadian time (CT) for DD, with white and black backgrounds denoting light and dark phases, or subjective day and night, respectively. **(C)** Dual-choice landing assay with *timeless* mutant females offered with a human-worn sleeve versus a clean one. The top panel shows preference index; asterisks indicate significant deviations from random choice (Binomial Exact Test, *p* < 0.05). The bottom panel shows landing proportions. **(D)** Behavioral preference of *timeless* mutants at ZT 0-2 (light blue) or ZT 10-12 (orange) under light-dark cycle, tested with artificial blends mimicking 6 AM (light blue), 6 PM (orange), or control (gray) odor profiles. Top: preference indices; bottom: landing proportions. **(E)** Behavioral preference of *timeless* mutants at CT 0-2 (light blue) or CT 10-12 (orange) under constant darkness, tested with artificial blends mimicking 6 AM (light blue), 6 PM (orange), or control (gray) odor profiles. Top: preference indices; bottom: landing proportions. Asterisks in top panels C, D and E denote statistically significant preference relative to random choice (Binomial Exact Test, *p* < 0.05). Asterisks in the bottom panels D and E indicate a significant difference with the positive control in C (Tukey post hoc test, * *p* < 0.05; ** *p* < 0.01; *** *p* < 0.001)

Our novel *timeless* knockout line allowed us to then define the role of the circadian clock in the phase-aligned attraction of mosquitoes to the host blends. When repeating the cage landing assays using these mutant, if light cues were sufficient to drive the time-dependent preferences, one would expect the *timeless* mutants to behave similarly to the wild-type mosquitoes under light-dark conditions.

Under both LD and constant darkness (DD) conditions, *timeless* knockout mosquitoes did not maintain the blend preferences observed in wild-type mosquitoes (Fig. 3C,D). Similar to the wild type (Fig. 2C), mutant females showed a significant preference for the positive control host (PI =0.29 ± 0.05, Binomial Exact test: *p* = 0.006, n = 10). However, the interaction between time of day and light regime was a significant predictor of preference (GLM: *p* = 0.024), and unlike wild-type mosquitoes, *timeless* mutants showed no attraction to the artificial blends, and in some cases even avoidance (Fig. 3D). Although their overall level of activity was significantly higher than wild-types under LD conditions in the activity monitors (Tukey adjusted *p* = 0.006), all treatment showed a significant reduction in the proportion of mosquitoes making a choice compared to the positive control for the mutants (Tukey-adjusted *p* < 0.002), except for the PM blend tested in the evening under LD (Tukey-adjusted *p* = 0.998) and the AM blend tested in the morning under DD (Tukey-adjusted *p* = 0.084).

Under LD conditions (Fig. 3D), mosquitoes tested in the morning showed a significant aversion for the AM (PI = - 0.52 ± 0.06, *p* < 0.001, n = 10) and PM blends (PI = -0.57 ± 0.08, *p* = 0.003, n = 9), with very low landing activity (landing proportion = 10 ± 1.46 %, *p* = 0.011 and 6.51 ± 1.19 %, *p* < 0.001 for the AM and PM blends, respectively). In the evening, no significant preference was detected for the AM blend (PI = -0.09, Binomial Exact test: *p* = 0.583, n = 11) and the PM blend (PI = -0.11 ± 0.05, Binomial Exact test: *p* = 0.337, n = 11), although the landing proportion was comparable to the positive control (AM: 13.87 ± 1.77 %, *p* = 0.571; PM: 18.30 ± 1.76 %, *p* = 0.998).

Under constant darkness (Fig. 3E), *timeless* mutants tested in their subjective morning were indifferent to either blend (AM blend: PI = -0.26 ± 0.06, Binomial Exact test: *p* = 0.056, n = 21; PM blend: PI = -0.15 ± 0.06, *p* = 0.26, n =10), and the landing activity was low, yet not significantly different from the control for the PM blend (12.52 ± 1.46 %, *p* = 0.106). However, landing activity significantly reduced when the AM blend was tested (6.34 ± 0.78 %, *p* < 0.001). In the subjective evening, *timeless* mutants significantly avoided the AM blend (PI = -0.66 ± 0.06, *p* < 0.001, n = 10) and were marginally repelled by the PM blend (PI = -0.37 ± 0.08, *p* = 0.050, n = 12; Fig. 3E). The proportion of mosquitoes landing on either cup was low yet not significantly different from the control for the AM blend (11.48 ± 1.69 %, *p* = 0.084), but significantly reduced when the PM blend was tested (8.55 ± 1.45 %, *p* = 0.002).

Together, our data suggest that *timeless* is involved in a regulatory network that maintains the olfactory preference for the blends’ volatiles and is required for the light-dependent modulation of the response throughout the day.

### Behavioral preferences are correlated with central and peripheral transcriptome modulations

The time- and light-dependent effects on mosquitoes’ responses to the host blends (Fig. 2), are likely dependent on daily variations in either sensory toolkit (*i*.*e*., antennal set of odorant receptors) and/or central neuromodulatory processes (*i*.*e*., fluctuations in brain neurotransmitter expression). To shed light on the underlying molecular mechanisms supporting the observed rhythms, we performed bulk RNA-sequencing of the head of *Ae. aegypti* females maintained under 12:12 LD cycles. We collected 4 replicates at 4 times of day, capturing the morning and evening activity peaks (ZT 0–2 and ZT 10–12, respectively) and troughs (ZT 5–7 and ZT 18–20, respectively)(Fig. 4, Fig. S4). We identified 1876 differentially expressed genes (DEGs) among the four timepoints (Likelihood Ratio Test (LRT), adjusted *p*-value < 0.05; Fig. 4). When contrasting DEGs to a list of sensory and neuromodulatory genes collated from previous studies in the field (29, 44–47), we detected a time-dependent effect on neuromodulatory genes (14 % [23 genes out of 161] of genes of interest) but only a few genes related to sensory functions (2.8 % [11 genes out of 391] of genes of interest, Fig. 4C,D). The low number of sensory genes found to be time-dependent could be explained by the relatively low number of cells expressing these genes in the whole mosquito head.

**Fig. 4.**
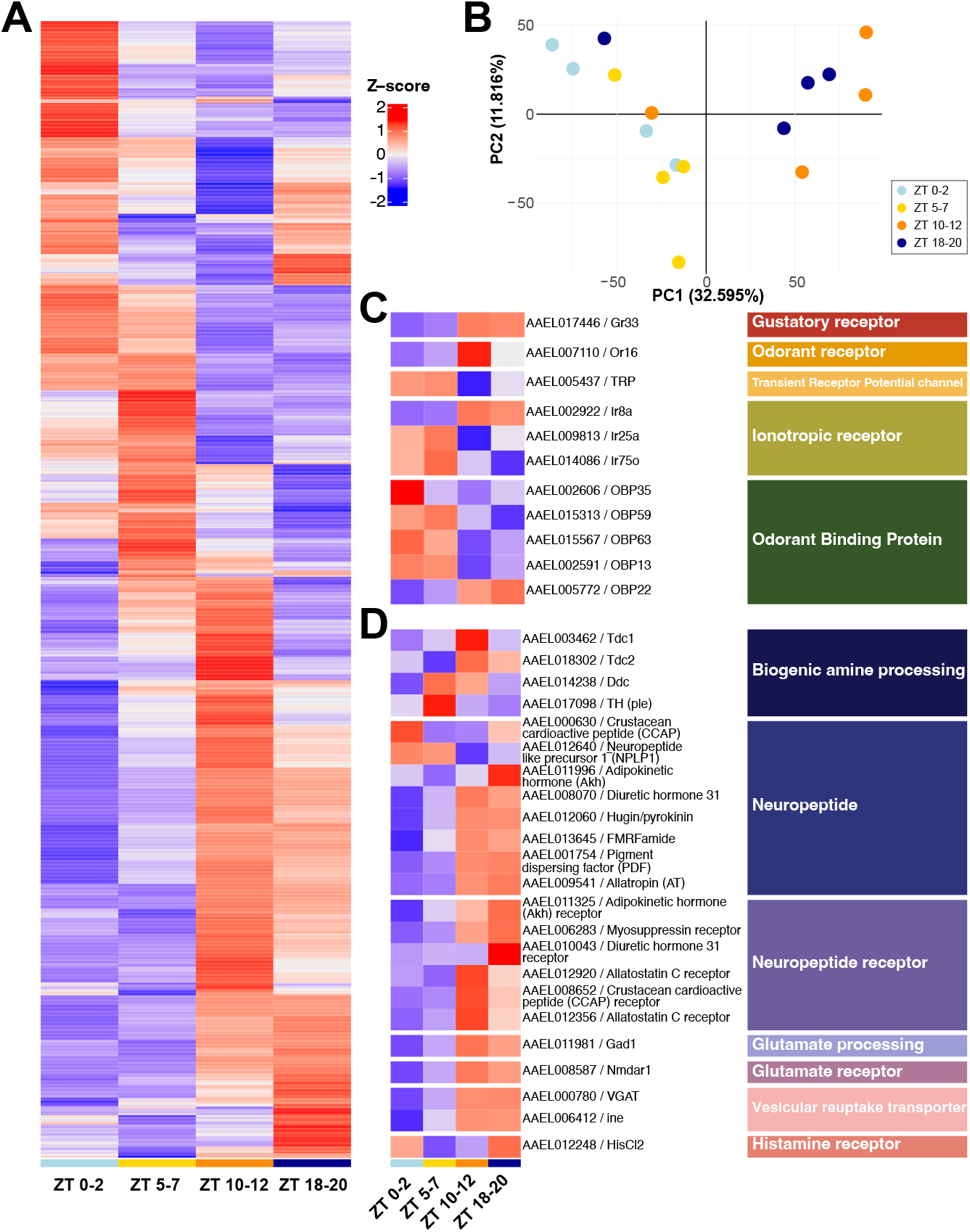
Gene expression daily rhythms in the head of *Ae. aegypti* females correlates with differences in neuronal integration throughout the day. **(A)** Heatmap of the *z*-score from averaged expression by timepoint of genes with time-of-the-day dependent expression (LRT, adjusted *p* < 0.05). **(B)** Principal Component Analysis (PCA) on head expressed genes at the four different timepoints. **(C-D)** Heatmap of the *z*-score from averaged expression by timepoint of genes with time-of-the-day dependent expression from a list of **(C)** sensory related genes of interest and **(D)** neuromodulatory genes of interest. For heatmap with individual replicates see Fig. S4.

To better understand how time-of-day specifically influences sensory gene expression, we then carried out bulk RNA-seq analyses on antennae collected every two hours throughout the day (*i*.*e*., 12 time points per day). Under a 12:12 LD regime (Fig. 5A, Fig. S5), 1558 time-of-day dependent DEGs were identified (LRT, adjusted *p*-value < 0.05; Fig. 5A). As expected, due to the dedicated sensory function of the antennae, only 492 DEGs were overlapping between head and antennae, with a higher proportion of sensory-related genes and fewer neuromodulatory genes being present in the antennae compared to the whole-head (Fig. 5A,E,F). We then compared time series in the head and antennae, and found 908 DEGs between the two types of samples (LRT, adjusted *p*-value < 0.05, |Log_2_FC| > 1.5) with 611 up-regulated genes in the antennae and 297 down-regulated (see Supplementary Materials), independently of the time of day.

**Fig. 5.**
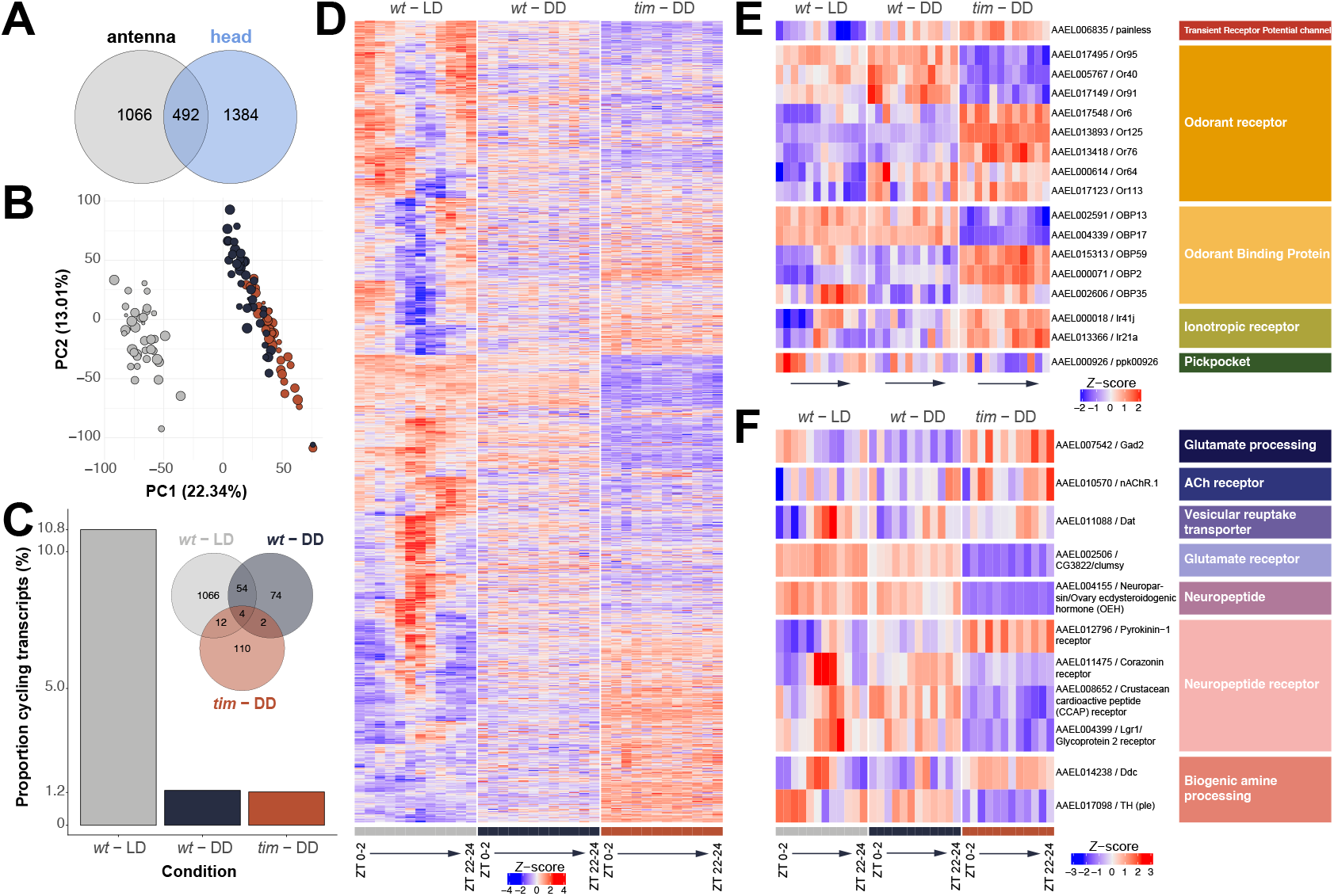
Light regime and *timeless* are essential for daily rhythms of expression of sensory genes in antennae of *Ae. aegypti* females. **(A)** Venn Diagram of the number of time-of-the-day dependent genes (LRT, adjusted *p* < 0.05) in head (blue) *versus* antennae (light grey) of *Ae. aegypti* females under light-dark cycles. **(B)** PCA on the full transcriptome of adult females antennae with the size of the dot representing the timepoint and the color the condition: wild-type in light-dark (light grey) and constant darkness (dark grey) conditions, and *timeless* mutants under constant darkness (brown). **(C)** Proportion of cycling genes (MetaCycle, *p*-value < 0.05, rAMP > 0.01) in the antennae of wild-type (*wt*) *Ae. aegypti* females in light-dark (light grey) and constant darkness (dark grey) conditions, and *timeless* mutants under constant darkness (brown). The Venn Diagram shows the raw number of cycling genes in each condition. **(D-F)** Heatmaps of the *z*-score from averaged expression by timepoint of **(D)** all genes with cycling expression throughout the day in the antennae of females *Ae. aegypti* (MetaCycle, *p*-value < 0.05, rAMP > 0.01) and subsets of genes of interest related to **(E)** the sensory system and **(F)** neuromodulation. For heatmap with individual replicates see Fig. S5.

Gene Ontology (GO) terms enrichment showed that DEGs between head and antennae were enriched in hormone and neuropeptide signaling pathways, heme and ion binding, fatty acid metabolism, and transporter activity GO terms (adjusted *p*-value < 0.05, see Supplementary Materials). Interestingly, some genes exhibited a peak of expression in the head at dusk (ZT10–12) suggesting time-specific neuromodulation (Fig. 4D): the aromatic amino acid decarboxylase (AAEL003462), an enhancer of split protein (AAEL004097), a calcium-binding protein (AAEL008844), two GTPases (AAEL011522 and AAEL003301) and two G-protein coupled receptors (GPCR) families: GPCR metabotropic glutamate family (*GPRMGL5*, AAEL009822) and GPCR galanin/allostatin family (*GPRALS3*, AAEL012920). Similarly, a set of genes exhibited a peak of expression at dusk in the antennae (Fig. S4) suggesting a tuning of signal transduction: inward-rectifying potassium channel (AAEL008932) and sodium-coupled cation-chloride co-transporter (*CCC1*, AAEL006180), a GPCR orphan receptor (*GPRNNB3*, AAEL001724), and an ionotropic glutamate receptor (AAEL013366). The expression of the ionotropic receptor *21a* (*Ir21a*, AAEL013366), which is involved in mosquitoes’ response to heat and thus associated with host-seeking (48), also peaked at dusk. All together these results suggest a time-dependent modulation of the sensory sensitivity at the peripheral level (*i*.*e*., antennae) and of the central sensory integration.

Given the significant effect of light cycles and the requirement for a functional molecular clock to sustain rhythms in olfactory preference, we next quantified the respective impact of these variables on rhythms in sensory gene expression. We collected antennae from both wild-type (*wt*) and *time-less* mutant (*tim*) adult females maintained under constant darkness conditions and using the same temporal sampling scheme (*i*.*e*., every two hours). The first striking difference between the three different conditions (*wt* LD, *wt* DD and *tim* DD) was the number of cycling genes: under light-dark conditions 10.8 % of the antennal transcriptome was cycling (MetaCycle, *p*-value < 0.05, rAMP > 0.01, Fig. 5B-D), while only 1.27 % and 1.22 % of the antennal transcriptome was cycling under constant darkness in the wild-types and *time-less* mutants, respectively (MetaCycle, *p*-value < 0.05, rAMP > 0.01, Fig. 5B-D). This suggests only a few genes in the antennal transcriptome are under true circadian regulation and highlights the dominant role of light-dark cycles in regulating the antennal transcriptome. Notably, only three genes (AAEL000071, *OBP2*; AAEL011088, *Dat*; AAEL011475, *corazonin receptor*) were cycling in wild-types under both LD and DD, but not in *timeless* mutants (Fig. 5E,F).

In terms of sensory genes, we identified one transient receptor potential channel (*painless*), involved in *Drosophila*’s mechanosensation, response to noxious heat (49), and time-dependent responses to colors (50), and a pickpocket channel (*Ppk00926*) involved in *Drosophila*’s mechanosensation (51), as being expressed in a time-dependent manner. Eight odorant receptor genes (*Or6*, indified to detect the plant-derived 4-chromanone (52), *Or40, Or64, Or76, Or91, Or95, Or113* and *Or125*), five odorant binding proteins (*OBP2, OBP13, OBP17, OBP35* and *OBP59*), and two ionotropic receptors (*Ir21a*, involved in heat and humidity sensing (48), and *Ir41j*) were also expressed in a time-dependent manner. The majority of these latter have not been deorphanized, making it difficult to link these rhythmic expression patterns with the observed behavioral responses to the blends.

### Gene expression networks underlie light and clock dependent effects

To determine whether daily light cycles and the circadian clock organize the antennal transcriptome beyond changes in individual genes (*i*.*e*., beyond differential gene expression), we examined how genes exhibited group-coordinated expression across the day. We used a Weighted Gene Co-expression Network Analysis (WGCNA) to characterize how groups of genes change in relation to the mosquito’s peak host-seeking window (ZT 10–12), allowing us to ask whether gene networks show coherent time-of-day structure under different experimental conditions (Fig. 6A). Under light–dark conditions (*wt* LD), the antennal transcriptome exhibited strong and highly ordered daily structure. Multiple gene groups showed coordinated increases or decreases in expression at specific times relative to the host-seeking peak (see *wt* LD -*plum1, greenyellow*, and *black* modules, Fig. 6A1). These coordinated patterns indicate that large sets of antennal genes are synchronously regulated in relation to behavioral timing. In contrast, the temporal organization was markedly weakened under constant darkness. In *wt* DD, only a small number of gene groups showed weak associations with time of day, and most genes no longer exhibited coordinated changes in expression across the circadian cycle (Fig. 6A2). This loss of structure indicates that light is a major driver of coordinated daily gene expression in the antenna. Disruption of the circadian clock further reduced temporal organization: *tim* DD mosquitoes exhibited the weakest alignment between gene expression groups and the host-seeking phase (Fig. 6A3). Together, these results demonstrate that both environmental light cues and an intact circadian clock are required to maintain a temporally structured antennal transcriptome.

**Fig. 6.**
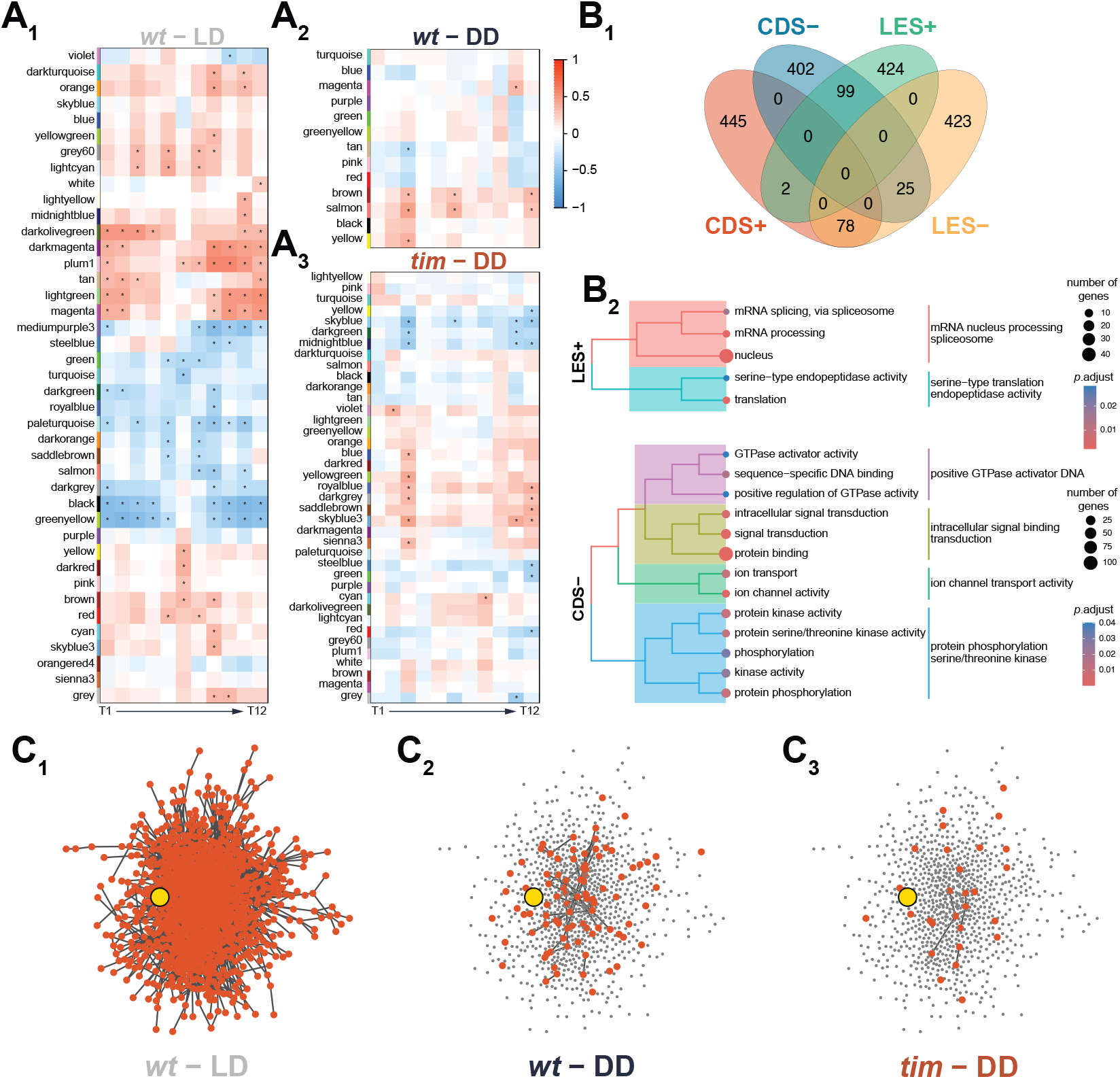
Network organization analyses revealed a distinct regulatory mechanism of light and circadian clock onto the mosquito antennal transcriptome. (**A_1−3_**)Heatmaps showing module eigengene–time correlations from independent WGCNA analyses performed separately for each experimental condition (*wt* LD, *wt* DD, and *tim* DD). Each row represents a co-expression module, and each column represents a time bin. The color gradient indicates the strength and direction of the correlation (red = positive; blue = negative). Asterisks denote statistically significant correlations. Because WGCNA was run independently for each condition, similar module names across panels do not indicate identical gene compositions. (**B_1_**) Venn diagram illustrating the classification of genes based on 5 % highest and lowest Light-Entrainment Scores (LES) and Clock-Dependence Scores (CDS). LES quantifies how a gene’s co-expression module membership shifts when light cues are removed, whereas CDS reflects changes in module membership when circadian clock function is disrupted. The top and bottom score percentiles identify genes most and least sensitive across experimental contexts. (**B_2_**) Gene Ontology (GO) term enrichment analysis for gene sets defined by LES and CDS categories. Terms summarize broad functional themes associated with light- and clock-sensitive changes in module membership. Circle size indicates the number of genes annotated to each term; color denotes adjusted significance levels. (**C_1_ − 3**) Protein–protein interaction (PPI) network projection of the *timeless*-associated co-expression module across experimental conditions. Genes from the module are mapped onto a fixed reference PPI framework, with TIM (protein from *timeless*) highlighted in yellow. Shown PPI edges and node color reflect their presence within the module containing *timeless* for each condition (*wt* LD, *wt* DD and *tim* DD). This visualization provides a framework for evaluating how the interaction landscape surrounding *timeless* is organized under different experimental contexts.

To formally assess which gene groups showed statistically robust time-of-day rhythmicity, we used a module-level analysis of expression trajectories across time (referred to as the Module Eigengene Trajectory Analysis or META), and identified 15 rhythmic gene groups in *wt* LD (FDR < 0.05), each exhibiting clear phase-dependent fluctuations in expression across the day (Fig. S6). In contrast, no rhythmic gene groups were detected in either *wt* DD or *tim* DD. Among the gene groups detected in *wt* LD, three exhibited exceptionally strong rhythmicity (FDR < 0.0001, η^2^ > 0.90), with circadian phase explaining more than 90 % of their variance in expression profiles. These included the previously reported *wt* LD *greenyellow* and *wt* LD *black* gene groups (Fig. 6A1). The strength and consistency of these rhythms suggest that a subset of antennal gene networks is tightly coupled to behavioral timing under natural light conditions.

To disentangle how light cycles and the circadian clock differentially shape the antennal gene network organization, we examined how strongly genes retained their network associations when either cue was disrupted. We did this by quantifying two complementary measures: a Light-Entrainment Score (LES), which reflects how much a gene’s network associations change when light–dark cycles are removed (*i*.*e*., from *wt* LD to *wt* DD), and a Clock-Dependence Score (CDS), which reflects how much those associations change when clock function is disrupted (*i*.*e*., from *wt* DD to *tim* DD). We selected the top and bottom 5 % of genes for each score (*i*.*e*., most and leasts impacted [*i*.*e*., most independent] genes, Fig. 6B) and performed GO term enrichment analysis. Genes with high LES values, those most affected by the absence of light cues, were enriched for transcriptional and post-transcriptional processes, including mRNA processing and splicing (Fig. 6B and Supplementary Material). This pattern indicates that light plays a dominant role in organizing nuclear and regulatory components of the antennal transcriptome. In contrast, genes least affected by clock disruption (low CDS values) were enriched for functions related to signal transduction, ion transport, GTPase activity, and kinase activity, suggesting that core sensory signaling pathways remain comparatively stable despite disruption of circadian timing (Fig. 6B and Supplementary Material). Together, these patterns indicate that light-driven and clock-dependent regulation act through distinct but complementary functional routes within the antenna.

Among genes implicated in maintaining temporal network organization, *timeless* illustrates how its position within the network, in addition to its own rhythmic expression, can drive temporal coordination. Although *timeless* itself is not part of a gene group classified as rhythmically oscillating (*wt* LD *turquoise*), it resides within a large, cohesive gene network that shows a pronounced decrease in expression around dusk (ZT12–14), coinciding with the peak and subsequent decline of host-seeking behavior. This network was enriched for pathways involved in autophagy, endocytic trafficking, and key signaling cascades (including mTOR, FoxO, MAPK, and phototransduction).

To examine how this network depends on light and clock function, we assessed the connectivity of *timeless*-associated genes within known protein–protein interaction frameworks (Fig. 6C). Under LD conditions, *timeless* (highlighted in yellow) is embedded in a large and densely connected interaction network consisting of 907 genes. In constant darkness (*wt* DD) and in clock-deficient mutants (*tim* DD), this connectivity collapses, with only 9.92 % and 3.20 % of the original interactions with *timeless* retained, respectively (Fig. 6C). This progressive loss of network architecture highlights the hierarchical importance of light cycles and proper TIME-LESS function in stabilizing temporally structured gene networks in the antenna.

Overall, these results indicate that *timeless* is embedded in a dusk-responsive gene network that is stable under light–dark conditions but reorganized in constant darkness (*wt* DD) and in *timeless*-deficient mosquitoes, suggesting a dusk-specific role for *timeless*, while other rhythmic gene networks coordinate a broader time-of-day-dependent sensory modulation.

## Discussion

Our findings show that mosquito–host interactions are temporally dynamic rather than static and shaped by both environmental light cues and the mosquito circadian clock, which align with subtle, yet consistent daily oscillations in human body odor. *Aedes aegypti* females displayed marked time-of-day differences in odor-driven activation: simplified synthetic blends increased behavioral responsiveness (14), but activation was strongest when the PM blend was presented during the mosquitoes’ evening activity phase. This indicates that host odor cues interact with the internal circadian state to gate olfactory-driven behavior.

Human body odor profiles varied strongly across individuals but only modestly across time of day. Nonetheless, several key volatiles, including nonanal, octanal, heptanal, and sulcatone, all known host selection and attraction markers (28, 32, 35, 53, 54), showed reproducible diel fluctuations. Even minor ratio-based changes in these compounds can influence mosquito attraction, consistent with evidence that the relative proportions of select volatiles, rather than total odor intensity, determine host preference (35, 37). Although the physiological basis for these odor rhythms is unknown, they likely reflect integrated changes in time awake, host metabolism, skin gland secretion, and microbial activity. These results indicate that mosquitoes can exploit these subtle temporal odor signatures, particularly during their natural foraging window.

Time-of-day differences in behavioral activation were mirrored by extensive circadian modulation of mosquito sen-sory and neuromodulatory pathways. Head transcriptomes showed dusk-enriched expression of genes involved in neurotransmitter synthesis (*e*.*g*., dopamine and histamine), calcium signaling, and both allatostatin C and mGluR pathways—suggesting central modulation of sensory gain, feeding state, and immune readiness prior to blood feeding (55– 57). At the periphery, ∼ 11 % of antennal transcripts cycled under light–dark conditions, including sensory receptors and ion channels peaking before dusk (Fig. 5C). Similar proportions of rhythmic gene expression were observed in the whole heads of *Ae. aegypti* (∼ 7 %, (22)) and *An. gambiae* (∼ 9-20 %, (58)), but higher than reported in the body of *An. gambiae* (∼ 4.5 %, (59)) and lower than in the salivary glands of *An. stephensi* (∼ 27 %, (60)). These rhythms are consistent with enhanced olfactory sensitivity near the active period and align with the report of dusk-peaking sensory gene expression in other mosquitoes (21). Genes supporting neuronal homeostasis and membrane repolarization (*e*.*g*., *Kir, CCC1*) also peaked at dusk, suggesting a lowered refractory period and higher excitability in antennal neurons (61). In addition, cycling expression of dopamine N-acetyltransferase (*Dat*, AAEL011088; Fig. 5E) suggests an inhibition of the dopamine pathway at dusk, associated with an increase in host-seeking activity in adult females *Ae. albopictus* (62). Finally, female mosquitoes up-regulated the GPCR orphan receptor (*GPRNNB3*, AAEL001724, ortholog of *GPCR143*) in their antennae at dusk. The GPCR orphan receptor is involved in stress response in moths, stimulating the transcription of heat shock proteins (63), proteins secreted by mosquitoes pre-bloodmeal (64, 65).

In constant darkness, mosquitoes lost blend-specific preferences and responded similarly to both AM and PM blends. These results suggest that while the endogenous clock contributes to olfactory responsiveness, external light cues are required to maintain phase-specific odor preference. This flattening of behavioral rhythms parallels the collapse of transcriptional oscillations in the antenna, where only ∼ 1 % of transcripts remained rhythmic under DD conditions (Fig. 5C). This result contrasts with previous reports of cycling gene expression in the whole head and body of various mosquito species for which the change in light regime reduces by a third the amount of cycling genes (22, 58, 59). Moreover, a recent study documented an increase of cycling genes in the salivary glands of *An. stephensi* under constant darkness, from ∼ 27 % in LD to ∼ 49 % of the whole salivary gland transcriptome (60). These studies (including this one) used similar algorithms to detect rhythmicity in gene expression and results thus reflect a biological phenomenon rather than an analytical artifact. We hypothesize that under constant environmental conditions, there is a benefit to lose time-dependent sensory tuning in the antennae, *i*.*e*., being able at all times to detect a variety of odors triggering behavioral responses (14). In contrast, oxidative stress from the ingestion of blood requires the secretion of a specific set of enzymes in anticipation of the blood meal. These enzymes are released by the salivary glands before the activity peak, when female mosquitoes are the more likely to find a host (66–68).

The *timeless* knockout line confirmed a critical role for the circadian clock in organizing these temporal behaviors. Although mutants retained general olfactory function, they lost the time-of-day–specific attraction seen in wild types under LD. The PM blend no longer elicited evening-specific attraction, and both blends were avoided during the early day. Under constant darkness, *timeless* mutants further failed to show any attraction to either blend. Thus, both light input and a functional circadian clock (via *timeless*) are required to maintain phase-aligned olfactory preference. At the molecular level under DD, not only did the mutant antennal transcriptome exhibit a similar amount of cycling genes compared to the *wt* DD group (Fig. 5C), but also a different level of expression of genes (Fig. 5D-F). Inversion in expression levels of sensory genes can explain the observed repulsion of the mutant to both blends in the morning, either due to under-activation of the over-expressed receptors or saturation of the under-expressed receptor proteins (Fig. 5E). Altogether, these results suggest a light-driven effect of antennal sensory system tuning by a cyclic expression of receptors, and a circadian-driven effect of the level of expression *per se* of sensory receptors in the antennae.

Network-level analysis of antennal transcriptomes revealed that temporal olfactory plasticity arises not from isolated rhythmic genes but from coordinated, phased reorganization of gene modules. Fifteen of 41 co-expression modules were rhythmic under LD, peaking near ZT 10–12, when host-seeking is maximal. In DD and in *timeless* mutants, these rhythmic modules lost temporal structure despite preserved network topology, indicating that light cues and the circadian clock maintain the modular timing that underlies sensory readiness. Genes most deeply embedded within rhythmic modules were the most sensitive to the loss of photic input or clock function, reflecting a regulatory hierarchy in temporal transcriptome organization (69). Furthermore, LES and CDS analysis associated with projected PPI analyses in the three different conditions revealed contrasting regulatory mechanisms of the antennal transcriptome by *timeless* and light cues. Similarly to what was observed in plants (70), light might impact differential splicing in mosquito antennae, a mechanism known to regulate insect immune specificity (71). In addition, light activation of *cryptochrome* in the antennal support cells has been shown to be essential for odor responses, especially for aversive responses, in flies (72). Given the interplay between CRYPTOCHROME and TIME-LESS proteins (73, 74), we hypothesized that *timeless* regulates the level of expression of sensory genes, while light triggers the time-dependent cycling of sensory genes and modulates the time-response specificity.

Together, these findings uncover a previously unrecognized temporal dimension of host seeking in mosquitoes. Human odor emissions, mosquito sensory physiology, and circadian state are phase-aligned in a way that enhances foraging efficiency during the evening activity window. Future work will determine whether temporal host cues may serve as external *Zeitgebers*, contributing to entrainment dynamics between vector and host, a largely unexplored axis of vector–host coevolution. This temporal coordination suggests new opportunities for chronobiology–informed vector control, including interventions that disrupt or shift the alignment between host scent rhythms and mosquito sensory peaks.

## Conclusions

Our results demonstrate that mosquito host seeking is shaped not simply by odor identity, but by the temporal alignment between host scent rhythms and the mosquito’s internal circadian state. By resolving how light cues, clock genes, and phased sensory gene networks converge to tune olfactory responsiveness, this work establishes time of day as a fundamental axis of mosquito–host interactions. Because blood feeding is the gateway to pathogen acquisition and transmission, such temporal tuning of host attraction is likely to structure when mosquitoes are most competent vectors, biasing transmission toward specific windows of host availability and vulnerability. Disrupting this alignment by targeting sensory gain, clock function, or environmental timing cues could therefore reduce mosquito–host contact without eliminating host cues themselves. More broadly, our findings reveal a previously unrecognized temporal dimension of vector–host coadaptation, in which rhythmic physiology on both sides shapes the ecology of disease transmission and exposes opportunities for time-based intervention.

## ACKNOWLEDGEMENTS

The authors thank Marium Khawaja, Dana Hamad, and Lily Smith for assistance with video analysis and cage-landing assays, and Seyed Jalil Pasha (JP) Mirlohi for assistance with maintaining the mosquito lines used in this study. The authors acknowledge the Advanced Research Computing (ARC) services at Virginia Tech for providing computational resources and technical support that have contributed to the results reported within this paper (URL: https://arc.vt.edu/). The following reagents were obtained through BEI Resources, NIAID, NIH: *Aedes aegypti*, Strain LVP-IB12, Eggs, MRA-735, and Strain ROCK, MRA-734, both contributed by David W. Severson. This work received direct support from the National Institute of Allergy and Infectious Diseases (NIAID) of the National Institutes of Health under Awards R01AI155785 (to C.V.), R21AI166633 (to J.B.B. and C.V.), which investigate the rhythms in mosquito olfactory processes and the mechanisms underlying mosquito sleep, respectively, and partial support from Award R01AI148551 (to J.B.B. for shared incubator space), as well as from the National Institute of Food and Agriculture, U.S. Department of Agriculture, through Hatch project VA-160212 (to C.V.).

## Methods

### A. Mosquito Rearing

Female *Ae. aegypti* Liverpool (LVP-IB12, MRA-735) and Rockefeller strains (ROCK, MR-734, MR4, ATCC®, Manassas, VA, USA) were used for behavioral experiments. Only the Liverpool strain was used for transcriptomic analyses and for establishing the *timeless* knockout line. Larvae were raised in a 26 x 35 x 4 cm covered tray filled with ∼ 1 cm of deionized water, at a density of approximately 200 larvae per tray. Mosquitoes were maintained under light:dark (LD) cycles of 12 h:12 h at 26 °C and 70 ± 10 % humidity. Larvae were fed a daily diet of Hikari Tropic First Bites (Petco, San Diego, CA, USA). Pupae were isolated on the day of pupation and placed into mosquito breeding containers (BioQuip, Rancho Dominguez, CA, USA). Mosquitoes emerged in the breeding containers and had unrestricted access to 10 % sucrose solution unless otherwise required by the experiment.

### B. Activity assay

Mosquito activity was measured using activity monitors that consist of 32 openings, with three sets of infrared emitters and detectors per opening (TriKinetics LAM25, Waltham, MA, USA). Each mosquito was placed into a clear cylindrical glass tube with mesh covering one end and a custom-made cotton plug with access to 10 % sucrose solution on the other end (See (14, 75, 76) for details). Tubes were then placed into the openings of the monitor so that the infrared beams bisected each tube in the middle. Activity monitors were placed inside a temperature-controlled chamber (model I-36VL, Percival Scientific, Perry, Iowa, USA), maintaining a temperature of 25 ± 1 °C and 40 ± 10 % relative humidity (RH) throughout the experiments. Mosquitoes were maintained under a 12-12 h light-dark cycle for 5 days with a 1 h long soft transition between light and dark to mimic sunrise (ZT 23:30-00:30) and sunset (ZT 11:30-12:30), *i*.*e*., gradual dawn and dusk transitions. At the beginning of the sixth day, mosquitoes were kept under constant darkness for 5 additional days. Daily activity was recorded as the number of beam crossings every minute using the DAMSystem3 Software (TriKinetics, Waltham, MA, USA). At the end of the ten days, the monitors were removed from the chamber, and mosquitoes were examined for survival. The data from tubes containing mosquitoes found dead at the end of the experiment (30 ± 10 % across experiments) were discarded from the analysis. To remove potential outliers, the 5 % of individuals with the least and highest number of beam breaks were excluded from the analysis.

### C. Unscented soap selection

Before sampling of human body odor, we conducted preliminary assays with five commercially available unscented soaps: Ivory (Mild & Gentle Body Wash, Fragrance-Free, Ivory Soap, Procter & Gamble, Cincinnati, Ohio, USA), Aveeno (Skin Relief Body Wash, Fragrance-Free, Kenvue, Summit, New Jersey, USA), Cetaphil (Ultra Gentle Body Wash, Fragrance-Free, Galderma, Fort Worth, Texas, USA), Native (Unscented Body Wash, Native, San Francisco, California, USA), and Dove (Hyper-Reactive Skin Balance, Fragrance-Free, Unilever, Englewood Cliffs, New Jersey, USA). The headspace profile of each soap was analyzed *via* Gas Chromatography -Mass Spectrometry and mosquitoes’ responses to the soaps were tested in dual choice landing assays (Fig. S1A). To ensure that soaps did not alter mosquitoes’ attraction for human hosts, we gave mosquitoes the choice between nylon stockings worn on the two arms of the same volunteer with one arm kept unwashed and the other arm washed with the test soap. Although none of the soaps, except Cetaphil (which produced a significant aversion; Binomial Exact test: *p* = 0.045), altered mosquitoes’ preference for human host, we selected the Native Unscented Body Wash for its low abundance in volatile and the absence of a preference or avoidance by mosquitoes (Fig. S1B). Anecdotally, we noted that while Dove produced a tendency to increase mosquitoes’ attraction to the host, this effect was not significant (Binomial Exact test, *p* = 0.124).

### D. Human body odor collection

Odor samples were collected from 12 adult participants (six males and six females) recruited from Virginia Tech (Blacksburg, VA) following previously published methods (35) (VT IRB #: 25-1196 and VT IRB #: 20-037). The volunteers included individuals from different ethnicities. Each participant provided at least four replicate samples at each of the following time points: 6 AM, 12 PM, 6 PM, and 12 AM.

The day before sampling, participants were required to use Native Unscented Body Wash (Native™, San Francisco, California, USA) instead of their regular body wash and to refrain from using perfume, deodorant, and other strongly scented products such as lotion.

Each participant’s left forearm was thoroughly cleansed for 30 seconds using deionized (DI) water and then dried using a disposable paper towel. Two strips of nylon sleeves, previously washed with unscented detergent and Tergazyme (Hanesbrands Inc., Winston-Salem, NC, USA/Ililily Inc., Irvine, CA, USA), each one inch in width, were applied to the forearm. The arm was then encased in aluminum foil to facilitate scent collection. This setup remained in place for one hour. During this period, participants remained active but tried to avoid activities that might significantly affect odor collection, such as bathing, cooking, swimming, etc. Following this period, the nylon sleeves were carefully removed, wrapped in aluminum foil, and placed in a ziplock bag for storage until processing and analysis.

Within 36 hours, nylon sleeves were placed into a nylon oven bag (Reynolds Kitchens, USA). The air within the bag was pulsed in using a diaphragm vacuum pump (400-1901, Barnant Co., Barrington, IL, USA) and directed through a headspace trap. This trap consisted of a Pasteur pipette loaded with 100 mg of Porapak Q powder (80-100 mech, Waters Corporation, Milford, MA, USA) and flanked by glass wool plugs (Restek, Belfonte, PA, USA). The air was purified through a charcoal filter before being reintroduced into the bag. The collection of headspace volatiles lasted 24 hours. These volatiles were subsequently extracted from the traps using 600 µL of hexane with a purity of 99 % (Sigma Aldrich, St. Louis, Mo, USA). The extracted samples were then stored in 2 mL borosilicate glass vials (VWR, Radnor, PA, USA) with Teflon-lined caps (VMR, Radnor, PA, USA) and maintained at a temperature of -80 °C. Analysis of the samples was conducted using a Gas Chromatograph-Mass Spectrometer (GCMS).

### E. Identification of rhythms in human scent chemistry

The liquid samples were injected into a GC-MS (GC: Trace 1310, MS: ISQ 7000, Thermo Fisher Scientific, Waltham, MA, USA) equipped with a 30-meter column (I.D. 0.25 mm, #36096-1420, Thermo Fisher Scientific, Waltham, MA, USA) and utilized helium as the carrier gas at a constant flow rate of 1 cc/min. The GC oven temperature was initially maintained at 45 °C for 3.75 minutes, then increased at a rate of 10 °C per minute until reaching 250 °C, and subsequently held constant at 250 °C for 10 minutes.

The Cobra algorithm within Chromeleon 7 software (Thermo Fisher Scientific, Waltham, MA, USA) was used to perform the integration of chromatographic peaks. Initial compound identification was conducted using the online NIST library, configuring the identification parameters to a normal search type with a minimum similarity index (SI) threshold of 600.

To facilitate quantification and normalization, 100 ng of heptyl acetate (HA) was added to each sample via pipette as an internal standard, enabling the conversion of peak areas into chemical quantities. A standard curve was prepared by serially diluting HA (1, 50, 100, 250, 500, and 1000 ng; CAS #112-06-1, Sigma-Aldrich, Saint-Louis, MO, USA) in hexane (Sigma Aldrich, St. Louis, Mo, USA).

To standardize the samples and remove additional contaminants, clean (*i*.*e*., washed), unused nylon sleeves were employed as negative controls following normalization with internal standards. The abundances of chemicals detected in these negative controls were subtracted from the analysis. Furthermore, hexane and inorganic chemicals were initially discarded to reduce noise from a large number of chemicals and focus on substances of interest. Chemicals accounting for less than 1 % of the total abundance in each sample were also excluded, as were chemicals not present in samples from at least seven participants. Chemicals previously reported to play a role in mosquito-human interactions were retained regardless of these criteria.

### F. Formulation of artificial human odor blends

Artificial odor blends were formulated to replicate the volatile profiles characteristic of human odor at either 6 AM or 6 PM. Each blend includes the same 12 chemicals, selected based on differential enrichment across time points, known relevance to mosquito olfaction, and accessibility through commercial suppliers. Raoult’s law was used to determine the partial vapor pressure contribution of each compound by adjusting their respective liquid-phase mole fractions, similarly to: (35, 37). Mole fractions were adjusted to achieve a total vapor pressure of 0.12 torr for each blend (*i*.*e*., the blends did not differ in overall intensity). Mineral oil was used as the solvent to prepare mixtures. Blends were freshly prepared in bulk, aliquoted into 1.5 mL centrifuge tubes to prevent volatilization, and stored at -20 °C until use. On the day of behavioral assays, aliquots were thawed and vortexed briefly before application onto filter papers.

The blends were prepared using the following chemicals: octanal (CAS 123-13-0, 99 %), 2-hexanone (CAS 591-78-6, 98%), heptanal (CAS 111-71-7, ≥ 95 %), 2-hexanol (CAS 626-93-7, 99 %), hexanal (CAS 66-25-1, 98 %), nonanoic acid (CAS112-05-0, ≥ 97 %), propenoic acid (acrylic aicd, CAS 79-10-7, 99 %), glutaric acid (CAS 110-94-1, 99 %), pentanal (CAS 110-62-3), 1-hexadecanol (CAS 36653-82-4, ≥ 99 %), nonanal (CAS 124-19-6, 95 %), and sulcatone (6-methyl-5-hepten-2-one, CAS 110-93-0, 99 %). All chemicals were purchased from Sigma-Aldrich, St Louis, MO, USA.

### G. Dual-choice assays

Behavioral assays were performed in a 30 x 30 x 30 cm cage (DP1000, BugDorm, Taiwan). Approximately 40 female mosquitoes, 5-7 days post-emergence, were cold-anesthetized at 4 °C for 10-15 minutes and transferred to individual holding containers on the day before testing, where they were deprived of sugar until the assays. To examine how human odor timing influences mosquito preference, we compared landing responses to artificial blends replicating either 6 AM or 6 PM odor profiles against odor-free controls.

For control assays, nylon sleeves (either clean or worn by a human volunteer for 1 hr) were placed in black plastic cups covered with fabric mesh to prevent direct contact with the mosquitoes, and symmetrically positioned inside the cage (Fig. 2B). For artificial blend assays, one-quarter of a filter paper treated with 250 µL of the odor blend (or, for controls, treated with 250 µL of mineral oil, which is the solvent used to dilute the blends) was placed inside the mesh-covered cups.

Mosquitoes were maintained in a light box under a 12:12 h light-dark cycle (LD) and temperature and humidity conditions similar to the experimental room, prior to experiments. Assays under LD were performed at either ZT 0-2 or ZT 10-12. For constant darkness (DD) experiments, mosquitoes were first entrained under LD for three days and then kept in DD for an additional three days before testing. DD assays were carried out under red light (*i*.*e*., functional darkness) to avoid photic interference.

Female mosquitoes were released into the cage for 30 min, during which their behavior was video-recorded (C920, Logitech, Lausanne, Switzerland) to minimize disturbance from the experimenter. Landings on each cup were scored from the recordings. A preference index (PI) was calculated as the number of mosquito landings on the treatment cup minus the number of mosquito landings on the control cup divided by the total number of landings. A landing proportion was also determined as the fraction of total landings relative to the number of mosquitoes tested. To reduce positional bias, the placement of odor and control cups was randomized between replicates. All assays were performed under well-ventilated conditions at 23 ± 2 °C and 50 ± 5 % relative humidity.

### H. Silencing of *timeless*

To knockout the *timeless* gene (AAEL019461) in *Aedes aegypti*, three single-guide RNAs (sgRNA) (see full data repository for details) were designed and synthesized according to previously described (77), targeting the second exon of the transcript AAEL019461-RA, coding sequence in common to all transcript isoforms of *timeless*. Approximately 180 embryos collected from females *Ae. aegypti* (Liverpool strain) that produce maternally deposited *Cas9* (Exu_Cas9, (78)) were injected with a mix of the three sgRNAs at 100 ng/µl each (77, 79). G_0_ adults were mated with the LVP adults of the opposite sex. Mutants G_1_ were screened using a high resolution melt curve analysis (80) followed by a 4 % agarose gel electrophoresis (79) to detect mutants with large deletions. We focused on one mutant that showed the largest deletions by crossing heterozygous mutants to generate homozygous. Purity of the line was checked by PCRs (Primers Tu_2280 and Tu_2247, see Supplemental Tables) and Sanger sequencing was performed on amplified fragments to identify the final 91 bp deletion in the wild-type sequence of the gene (Fig. S3).

### I. Gene expression analysis

#### I.1. Mosquitoes’ synchronization

Mosquito pupae were collected and placed into mosquito breeding containers on the day of pupation, which were then enclosed within light-tight plastic containers (Rubbermaid Brute boxes, 71 x 44 x 38 cm) equipped with LED lighting to achieve rhythmic synchronization. The light cycle implemented was a 12-hour light:12-hour dark (LD) cycle, with *Zeitgeber* Time 0 (ZT 0) marking the onset of light and ZT 12 marking the onset of darkness. Mosquitoes were synchronized for at least 3 days before experimental treatment (either LD or DD) for 3 more days. All experiments were performed in four biological replicates, and under red light for night time intervals or constant darkness treatment.

#### I.2. Sample collection and RNA isolation

##### i). Whole head RNAseq

The experiment included four time intervals (ZT0-2, ZT5-7, ZT10-12, ZT18-20), during which sample collections were conducted. The ZT18-20 experimental groups were handled under red light, while the remaining experimental groups were completed under normal light conditions.

After circadian synchronization, adult female mosquitoes aged 4 to 7 days post-emergence were decapitated, and their heads were collected into 1.5 mL microcentrifuge tubes kept on ice. Each sample consisted of 14-30 mosquito heads, and each experimental set included four biological replicates. Total RNA was extracted from each sample using the RNeasy Mini Kit (Qiagen, Hilden, Germany) and following the manufacturer’s instructions.

##### ii). Antennal RNAseq

In the LD experimental group, mosquitoes were exposed to a 5 to 7 day LD cycle to synchronize their activity with the experimenter’s working hours. In all cases, tested mosquitoes were removed from the synchronization box only after the light phase had begun, and all experiments were completed within 2 hours. For the DD experimental groups, mosquitoes were initially exposed to an LD light cycle for at least 3 days, followed by a minimum of 3 days in constant darkness. Experiments involving mosquitoes from the DD groups were conducted under red light illumination in all instances.

After circadian synchronization, antennae were carefully dissected, as intact as possible, from the heads of adult female mosquitoes aged 5 to 7 days post-emergence and immediately placed into a 1.5 mL microcentrifuge tube kept on ice. For each sample, approximately 200 mosquitoes were dissected to obtain around 400 antennae. The samples were then stored at -80 °C until the RNA extraction step. Each test set contained three to four biological replicates. Antennae were mechanically homogenized using a disposable DNase/RNase-free pestle (Cole-Parmer, Chicago, IL, USA), and total RNA was extracted using the RNAqueous™-Micro Kit (Thermo Fisher Scientific, Waltham, MA, USA).

#### I.3. RNA sequencing and RNA-seq data processing and analysis

Concentration and RNA integrity of the samples were assessed using a Nanodrop spectrophotometer (Thermo Scientific, Massachusetts, USA) and running the samples onto a 1 % agarose gel electrophoresis. Samples were sent to Novogene (Novogene Corporation Inc., Beijing, China) for mRNA poly(A) enrichment library preparation and 150 bp paired-end sequencing on Illumina NovaSeq, targeting around 25 million reads per sample for antennae samples, and 20 million reads per sample for head samples.

#### I.4. Sequencing quality check and reads mapping to the reference genome

Quality of the sequencing for each sample was assessed under FastQC v. 0.12.1 (81) before trimming of remaining adaptors and 10 first bp with Cutadapt v. 5.0 (82). All reads were then mapped onto the lastly available reference genome for *Aedes aegypti* (LVP_AGWG, release 68) from VectorBase (83). Read alignment was carried out with the STAR version 2.6.1b algorithm (84) and gene abundance computed by RSEM 1.2.28 (85). All quality checks and mapping rates were summarized under MultiQC v. 1.28 (86). Mapping from *timeless* knock-out samples were indexed with samtools version 1.21 (87) and visualized under IGV 2.19.4 (88) to remove samples showing high heterozygosity contamination at the deletion sites.

#### I.5. Normalization, differential gene expression and cycling rhythmicity

All subsequent analyses were performed in *R* version 4.5.0 (*R* Core Team, 2025). Analyses were first performed on the head time series (∼ ZT) and antennae wild-type LD cycles time series (∼ ZT) before comparing head and antennae time series (∼ ZT+tissue) and finally the three antennae time series (∼ ZT + condition, considering *wt* LD, *wt* DD and *tim* DD as three independent conditions). First, low count genes (< 10 counts) were filtered out before normalization of the counts by library size, timepoint and condition (see above) using the DESeq2 pipeline version 1.48.0 (89).

Normalized counts were exported using the *rlog* transformation function for the same package. A likelihood ratio test (LRT) was performed to identify the effect of ZT time on the expression of genes in the head and antennae (wild-type under LD cycles). For the antennae dataset, rhythmicity of gene expression in the three different conditions for each gene was tested under MetaCycle 1.2.0 (90) combining JTK_CYCLE (91) and Long-Scargle (92) methods using the *meta2d* function of the package. Genes were considered as rhythmic if they fitted into the combined *p*-value of the two methods threshold (*p*-value < 0.05) and exhibited a relative amplitude through time of at least 1 % (rAMP > 0.01). Heatmaps were generated with the Complex Heatmap package 2.23.1 (93), and Gene Ontology (GO) terms enrichment analyses performed with clusterProfiler v. 4.17.0 package (94).

### J. Weighted Gene Co-Expression Network Analysis (WGCNA)

Weighted Gene Co-expression Network Analysis (WGCNA) was conducted on the antennae dataset separately for each treatment condition (*wt* LD, *wt* DD, and *tim* DD) to identify groups of co-expressed genes and characterize their temporal organization (69, 95). Expression matrices were pre-processed by log transformation, removing low-variance and sparsely detected genes, and retention of genes present across all three treatment conditions to ensure compatibility across networks. All networks were constructed using signed Pearson correlations, such that only positively correlated expression profiles contributed to module formation (95). This choice preserves the directional interpretation of co-expression, ensuring that genes grouped together exhibit coordinated increases or decreases in expression across time, rather than anti-phase behavior. The correlation matrix was raised to an empirically selected soft-thresholding power to approximate scale-free topology (69). The resulting adjacency matrix was converted into a topological overlap matrix (TOM), and genes were hierarchically clustered based on TOM dissimilarity (95).

Gene groups (also referred to as *modules*) were identified by clustering genes with similar expression patterns over time, with a minimum group size of 30 genes. Groups with highly similar overall expression profiles (correlation > 0.75) were subsequently merged to represent a single coordinated expression pattern (96). In wt DD, the gene expression data exhibited weak co-expression structure, yielding only a few gene groups with low internal coherence and no distinct *grey* group. As a result, genes were broadly assigned to these low-structure groups rather than remaining unassigned. Each group was summarized by its eigengene, representing the primary shared expression pattern of its member genes (96).

To ensure accurate interpretation of positive and negative correlations, we manually inspected representative expression trajectories for genes showing strong positive or negative correlations with module eigengenes, particularly in wt DD and *tim* DD. In these conditions, negative correlations often reflected flat or weakly varying expression rather than true downregulation, consistent with the overall loss of temporal structure in these networks (97, 98). Negative correlations were therefore interpreted as loss of coordinated rhythmic behavior rather than phase-opposed regulation.

### K. Temporal association and phase-specific analysis

To assess how gene groups varied across the day, we related each group’s overall expression pattern to two-hour time windows and evaluated how expression at each time point differed from ZT10–12, the period of mosquitoes’ peak host-seeking activity (21, 59). Statistical significance was determined using correlation analyses with false discovery rate (FDR) correction, allowing us to identify gene groups whose expression changed systematically at specific circadian phases relative to this behavioral reference. For clarity and consistency, gene groups are labeled by treatment condition and color (for example, *wt* LD *greenyellow, wt* DD *black*, or *tim* DD *violet*) rather than by eigengene identifiers. Because networks were constructed independently for each condition, this nomenclature avoids implying that similarly colored gene groups represent the same biological entities across treatments.

### L. Projection-based metrics for light and clock dependence

This network-level analysis was complemented by gene-level metrics to quantify how well individual genes maintain their network relationships when light cues or clock function are disrupted. The *wt* LD network was used as the reference because it displayed the strongest temporal organization and the most clearly defined co-expression structure. Using this reference allowed us to assess how much of the light–dark–organized network architecture persisted under constant darkness or clock disruption. For each gene, network association was quantified as module membership (kME), defined as the correlation between the gene’s expression profile and the expression pattern representing its assigned *wt* LD gene group (95). These same reference expression patterns were then evaluated in *wt* DD and *tim* DD, allowing us to determine how strongly each gene continued to track its original *wt* LD network association under each perturbation. A single, combined WGCNA across treatments was not performed, because the weak and poorly structured networks observed in *wt* DD and *tim* DD would mask the rhythmic organization present under *wt* LD conditions (95).

From these comparisons, two composite measures were derived. The Light-Entrainment Score (LES) measured how strongly a gene’s network association depends on light cues, based on the degree to which its *wt* LD network membership and connec-tivity weaken in *wt* DD (the change in |kME| from LD to *wt*-DD). The Clock-Dependence Score (CDS) evaluates the additional loss of network association that occurs when circadian clock function is disrupted, based on changes observed between *wt* DD and *tim* DD (the change in |kME| from wt-DD to *tim*-DD). Both scores were standardized prior to combination, such that higher values reflect greater disruption of the *wt* LD–defined network organization. To relate these network changes to biological function, Gene Ontology (GO) enrichment analysis was performed on genes showing the strongest and weakest dependence on light (LES+ and LES-) and clock function (CDS+ and CDS-), defined as the top and bottom 5 % of each score (99, 100). This approach links changes in network organization to functional gene categories, providing biological context for how light and the circadian clock shape antennal gene networks.

### M. Module Eigengene Trajectory Analysis (META)

To formally test whether gene groups display a reliable time-of-day structure across the circadian cycle, we employed the Module Eigengene Trajectory Analysis (META), a statistical framework that evaluates whether the overall expression pattern of a gene group changes systemically across time (96, 97). META assesses these changes relative to ZT10–12 (timepoint T6 in the full dataset), the period corresponding to peak host-seeking activity, allowing us to determine whether gene groups exhibit consistent, phase-dependent increases or decreases in expression across the day. For each gene group, we tested whether its overall expression pattern differed across time bins using a one-way analysis of variance (ANOVA), with time of day as the explanatory factor. Statistical significance was evaluated after correcting for multiple comparisons using the Benjamini–Hochberg procedure. In addition to significance, META quantifies effect size (η^2^), which represents the proportion of variation in a gene group’s expression that can be explained by circadian phase (101). Values approaching 1 suggest that module activity is nearly entirely driven by circadian phase, whereas values closer to 0 indicate a lack of temporal structure. Gene groups with η^2^ values greater than 0.90 were considered to exhibit exceptionally strong rhythmicity, indicating that their expression dynamics are dominated by time-of-day effects rather than random variation. By combining WGCNA-based identification of co-expressed gene groups with META-based statistical validation, this approach identifies gene groups that are not only associated with circadian phases but also robust, formally validated rhythmic organization across the day.

### N. Protein-protein interaction network projection analysis

In order to visualize the effect of light and *timeless* disruption on the antennal gene network organization, we recovered the protein-protein interaction (PPI) network of the WGCNA module containing the *timeless* gene using Cytoscape StringApp version 2.2.0 (102) in Cytoscape version 3.10.4 (103) with the *Aedes aegypti* database. Main network (907 nodes) including *timeless* was then imported in *R*, filtered out by condition for nodes and edges included in the network of *timeless* and plotted with ggraph package version 2.2.2 (104).

## Supplementary Figures

**Fig. S1.**
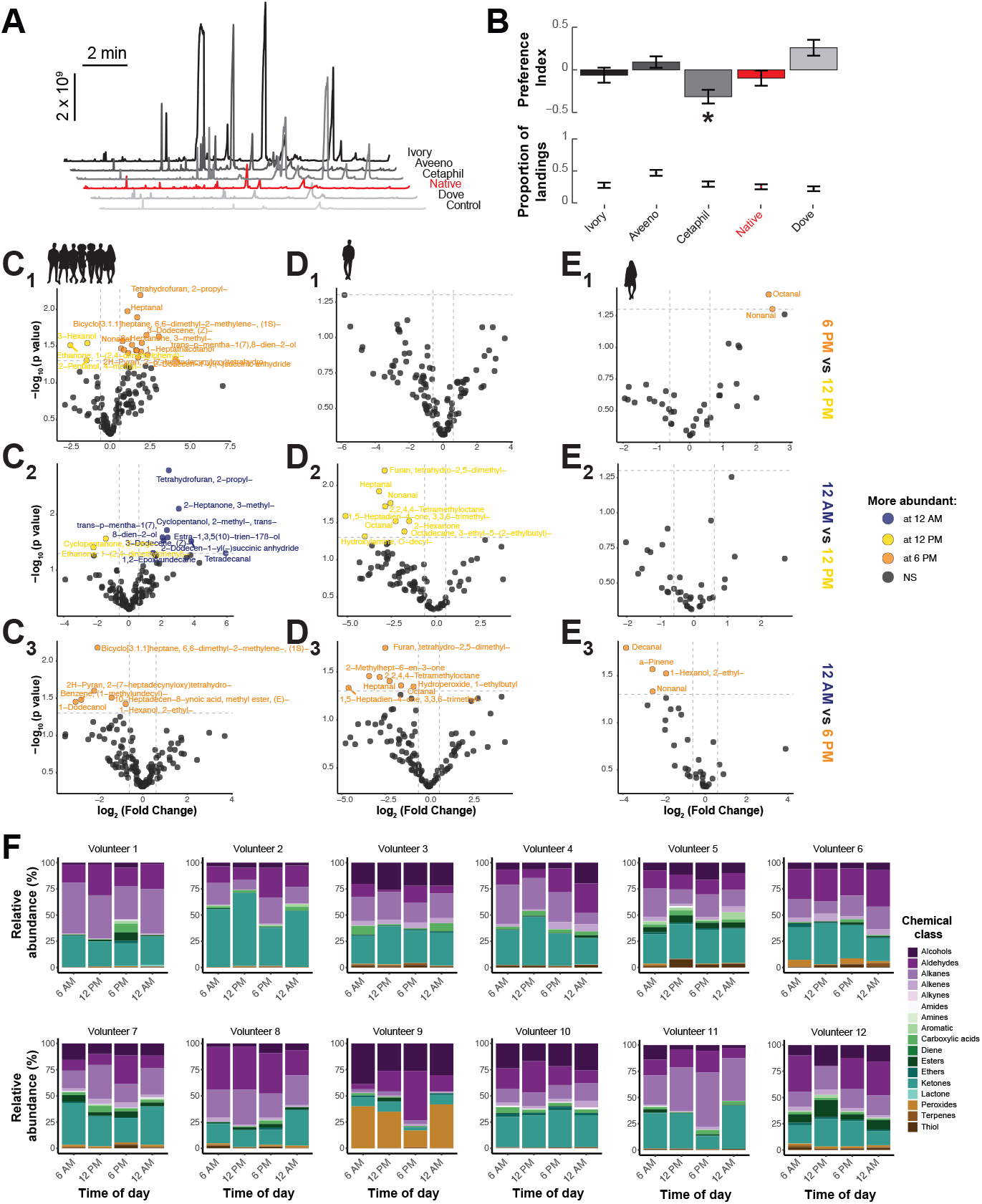
Daily chemical shifts in human body odor profiles. **(A)** Representative GC-MS chromatograms of headspace volatiles from five commercially available unscented soaps (Ivory, Aveeno, Cetaphil, Native, and Dove) and a negative control. The chromatogram for Native Unscented Body Wash is highlighted in red and selected for subsequent human odor sampling. **(B)** Preference index (top) and proportion of landings (bottom) in dual-choice landing assays in which mosquitoes chose between nylon sleeves worn on two arms of the same volunteer, with one arm washed with the indicated soap and the other unwashed. Cetaphil induced a significant aversive effect (asterisk: Exact Binomial test, *p* = 0.045), whereas the rest soap did not significantly alter attraction. (**C**_1_-**C**_3_, **D**_1_-**D**_3_, **E**_1_-**E**_3_) Volcano plots showing differential enriched compounds between sampling time points for the (**C**_1_-**C**_3_) combined multi-host dataset, (**D**_1_**-D**_3_) a representative male host and (**E**_1_**-E**_3_) a representative female host. Colored points highlighted compounds significantly more abundant at 12 AM (dark blue), 12 PM (yellow), or 6 PM (orange); gray points are not significantly different (log_2_(fold-change) > 0.6, adjusted *p* < 0.05). **(F)** Relative abundance of chemical classes in human body odor profiles across time of day for each individual. Stacked bar plots show the proportional contribution of chemical groups to the overall odor profile at 6 AM, 12 PM, 6 PM, and 12 AM.

**Fig. S2.**
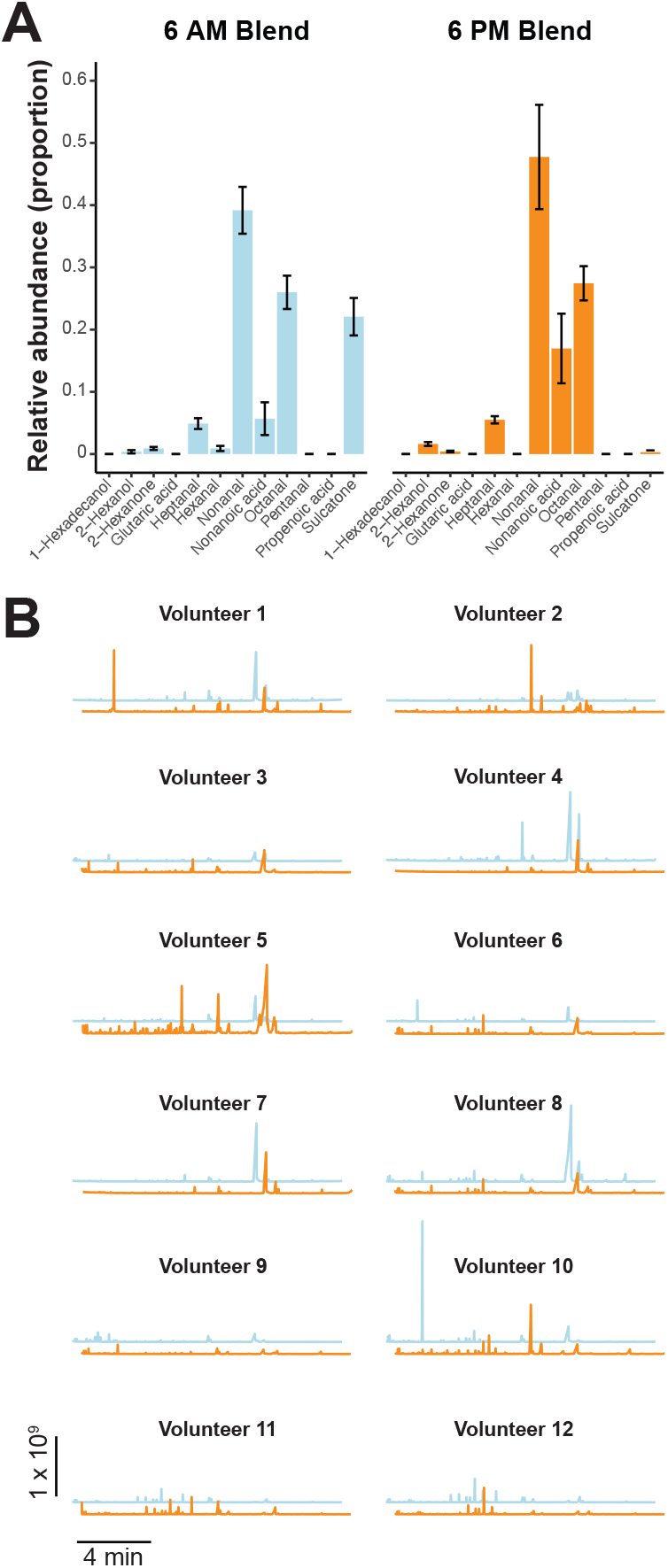
Artificial blend formulation and individual host variation in odor profiles. **(A)** Relative composition of the 6 AM (left, blue) and 6 PM (right, orange) artificial blends, as quantified by GC-MS analysis of dynamic headspace samples collected from 25 µL of each blend. Bars show the normalized contribution of each compound to the total blend. **(B)** Representative GC-MS chromatograms of human body odors collected at 6 AM (blue) and 6 PM (orange) from each of the 12 volunteers. The scale bars indicating the retention time (minutes) and the abundance (arbitrary units) apply to all panels.

**Fig. S3.**
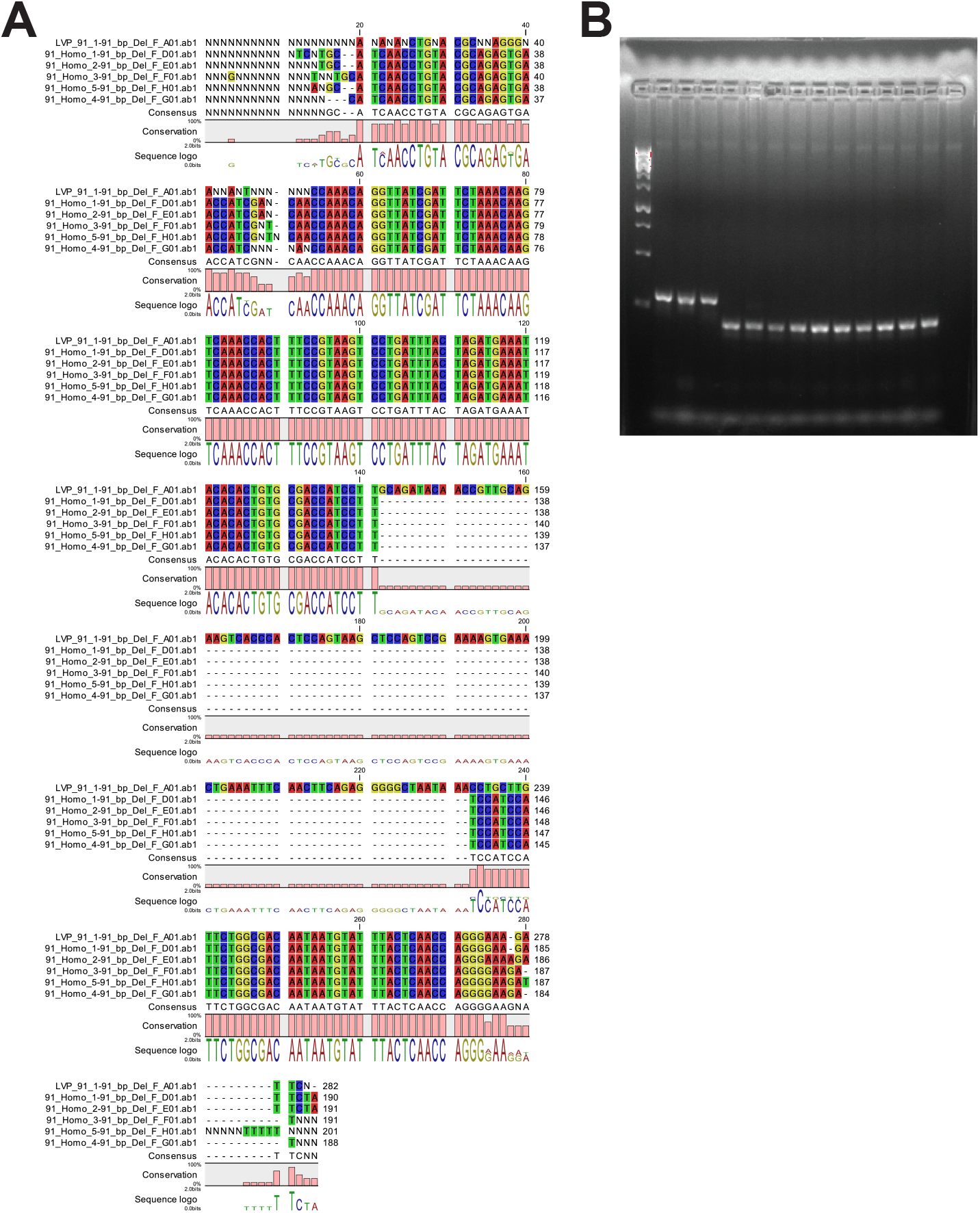
*Timeless* knockout and experimental validation. **(A)** Alignment of Sanger sequencing reads from the *timeless* 91-bp deletion line (5 bottom rows) compared with the wildtype reference sequence (top row). **(B)** Representative agarose gel showing diagnostic PCR products spanning the edited region in wild-type and *timeless* mutant mosquitoes. The mutant allele (lanes 5-14) yields a shorter amplicon compared with the wild-type fragment (lanes 2-4), confirming the presence of the 91-bp deletion.

**Fig. S4.**
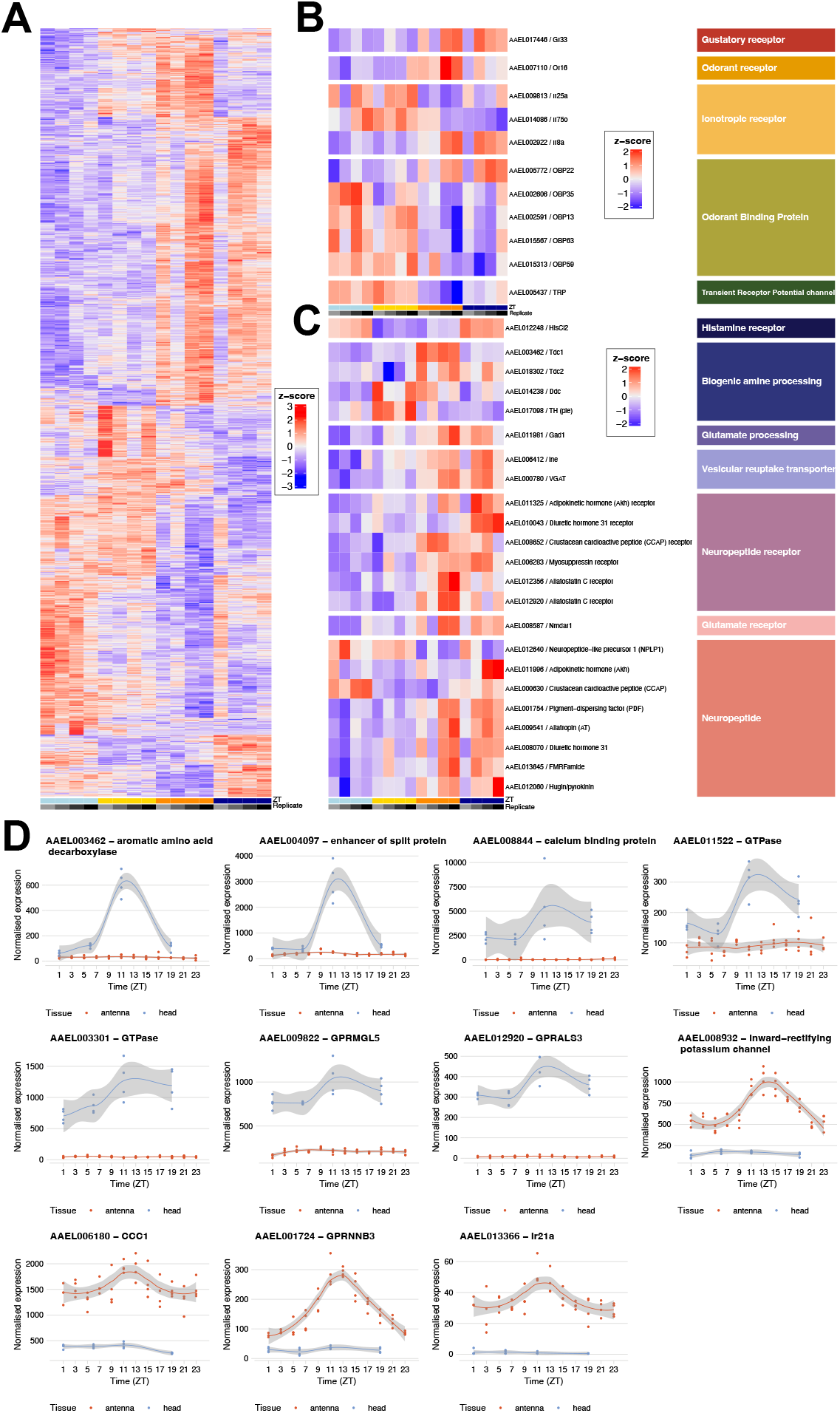
Gene expression daily rhythms in individual head samples of *Ae. aegypti* females. **A)** Heatmap of the *z*-scored expression level of each replicate as a function of time, for genes with time-of-the-day dependent expression (LRT, adjusted *p* < 0.05). **(B-C)** Heatmap of the *z*-score individual replicate expression by timepoint of genes with time-of-the-day dependent expression from a list of **(B)** sensory related genes of interest and **(C)** neuromodulator genes of interest. **(D)** Normalized expression of selected genes showing peak expression at dusk in either the head (blue lines) or the antennae samples (red lines). Solid lines indicate a loess fit on the measured data (points), and the shaded areas indicate the 95 % confidence interval around the mean.

**Fig. S5.**
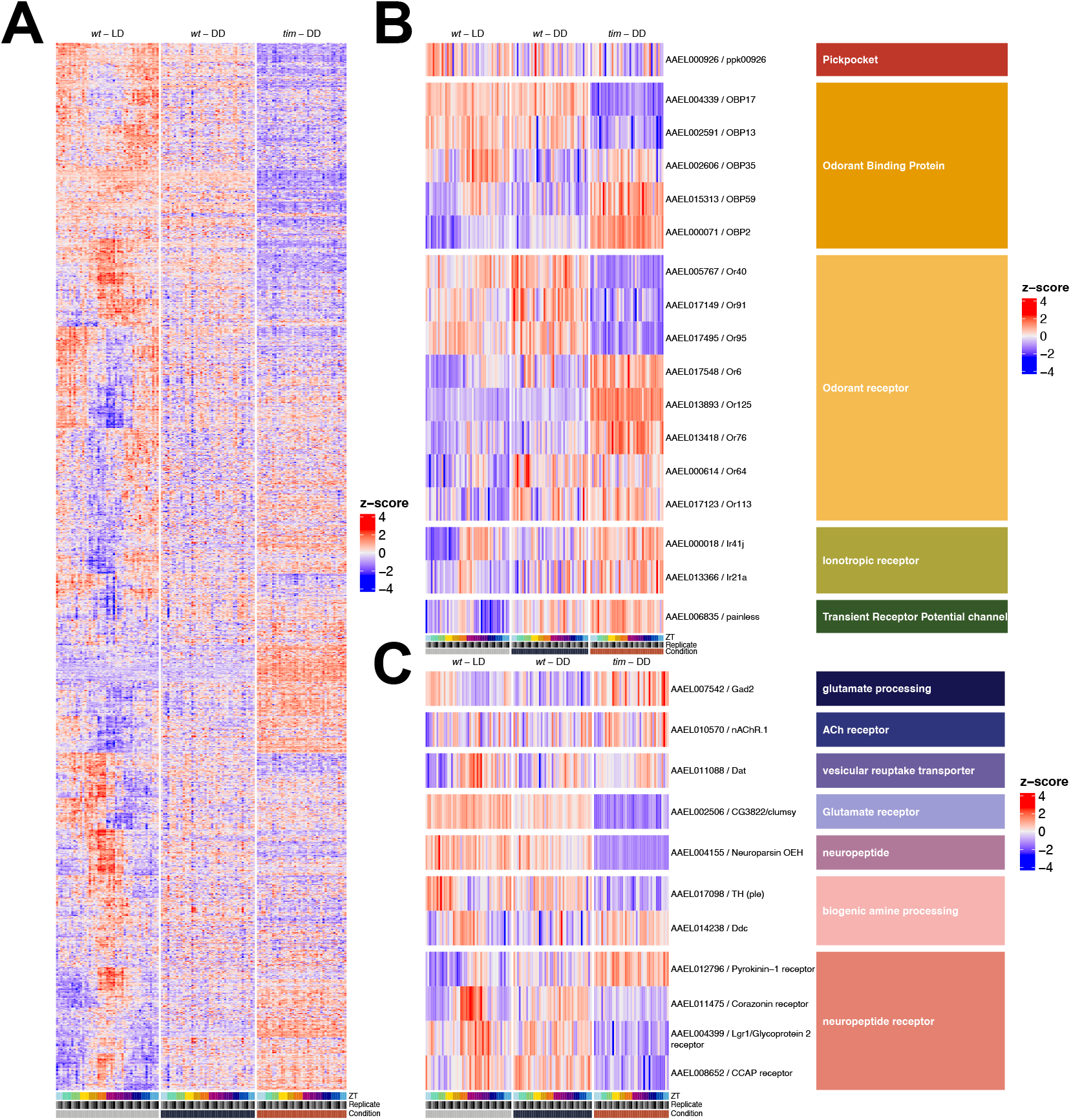
Gene expression daily rhythms in individual antennae samples of *Ae. aegypti* females. Heatmaps of the *z*-score from individual replicates’ expression by timepoint of **(A)** all genes with cycling expression throughout the day in the antennae of females *Ae. aegypti* (MetaCycle, *p*-value < 0.05, rAMP > 0.01) and subsets of genes of interest related to **(B)** the sensory system and **(C)** neuromodulation.

**Fig. S6.**
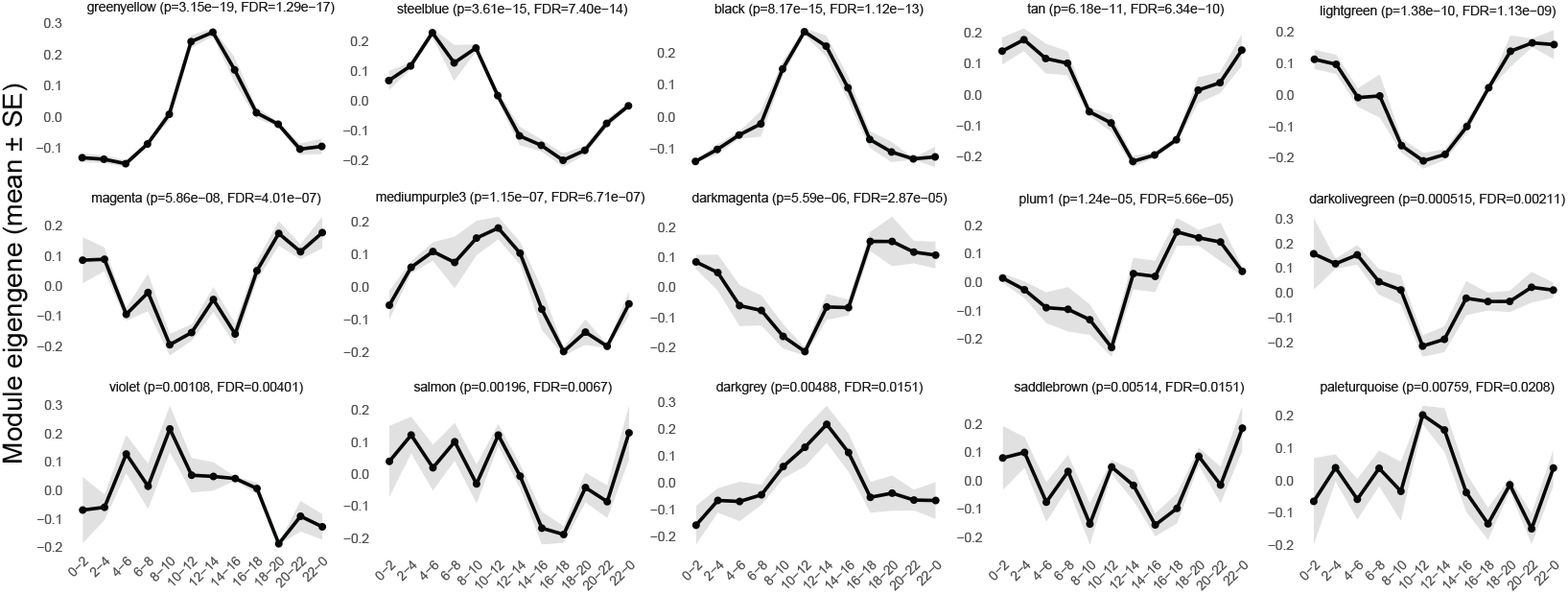
Time series of module eigengenes from the WGCNA. Normalized averaged expression level of the eigengene of each of the twelve modules showing significant time-dependency as a function of the RNA sample’s collection time. Shaded areas indicate the 95 % confidence interval around the mean. Statistical details are provided in the subtitle above each graph.

